# A novel subset of colonocytes targeted by *Citrobacter rodentium* elicits epithelial MHCII-restricted help from CD4 T cells

**DOI:** 10.1101/2023.05.03.539269

**Authors:** Carlene L. Zindl, C. Garrett Wilson, Awalpreet S. Chadha, Baiyi Cai, Stacey N. Harbour, Yoshiko Nagaoka-Kamata, Robin D. Hatton, Min Gao, David A. Figge, Casey T. Weaver

## Abstract

Interleukin (IL)-22 plays a non-redundant role in immune defense of the intestinal barrier^1–3^. We recently discovered an indispensable role for T cells, but not ILCs, in sustaining IL-22 signaling required for protection of colonic crypts against invasion during infection by the enteropathogen, *Citrobacter rodentium* (*C.r*)^4^. However, identification of the intestinal epithelial cell (IEC) subsets targeted by T cell-derived IL-22 and how T cell-derived IL-22 sustains activation in IECs are undefined. Here, we identify a novel subset of absorptive IECs in the mid-distal colon that are differentially targeted by *C.r* and are differentially responsive to IL-22 signaling. Importantly, MHCII expression by these colonocytes was required to elicit T cell-activated IL-22 signaling necessary to resist *C.r* invasion. Our findings explain the basis for the regionalization of the host response to *C.r* and demonstrate that epithelial cells must elicit MHCII-dependent help from IL-22–producing T cells to orchestrate immune protection in the intestines.

## Introduction

The arc of an infectious response is determined by the tissue site that is targeted by a pathogen and the host defenses mounted at that site. In vertebrates, which employ adaptive immunity to confer greater specificity and memory to host defense, interplay of the innate and adaptive immune systems is central to the anti-pathogen response and is coordinated with tissue-specific, non-immune cell populations to resist and defeat pathogen incursion, with specialization of immune response types predicated on the class of pathogen^5–8^. While host defense against phagocyte-resistant intracellular pathogens and extracellular parasites and helminths are dependent on type 1 and type 2 responses, respectively, host defense against extracellular bacteria is orchestrated by type 3 immunity, which employs innate and adaptive immune cells that are responsive to IL-23 and produce IL-17 family cytokines and the IL-10 family cytokine, IL-22^1, 2, 9^. These cytokines act on non-immune cells, including barrier epithelial cells, to bolster anti-microbial responses.

In some infections, lack of one of these cytokines is fatal^1, 9^. *Citrobacter rodentium* (*C.r*) is a murine enteropathogen that belongs to a family of genetically related Gram-negative bacteria that includes enterohaemorrhagic and enteropathogenic *E. coli* (EHEC and EPEC), all of which form attaching and effacing (A/E) lesions on the apical surface of intestinal epithelial cells (IEC) via a type III secretion system that delivers membrane-associated and intracellular virulence factors to the host^10–14^. IL-22 produced by both innate and adaptive immune cells is indispensable for host protection against *C.r*^1–4, 15^, acting on IECs to induce STAT3-mediated signaling^4, 16^ required to upregulate antimicrobial peptides (e.g., Reg3 and S100a family members)^1, 17, 18^, neutrophil-recruiting chemokines (e.g., Cxcl2 and Cxcl5)^19–21^ and mucin-related molecules (e.g., Fut2 and Muc1)^22, 23^ that protect the epithelial barrier. Production of IL-22 is spatially and temporally constrained to different innate and adaptive immune cells^1, 3, 4, 15^. Resident innate cells (type 3 innate lymphoid cells (ILC3)^2, 8, 15, 24–29^, natural killer (NK) cells^29–31^, NKT cells^32, 33^, γδ^+^ T cells^34, 35^, and neutrophils^36–38^ produce IL-22 in response to IL-23 and are dominated by ILC3s during the early phase of *C.r* infection, whereas CD4 T cells (Th17 and Th22)^3, 17, 39–43^ can act independently of IL-23^3, 9, 39, 44^ and are the dominant source of IL-22 late following their recruitment to the lamina propria (LP) of the distal colon^4, 45^. Accordingly, deficiency of IL-22 targeted to innate immune cells results in fatality early in *C.r* infection, whereas IL-22 deficiency targeted to T cells results in fatality late^4^.

A basis for the subdivision of labor among IL-22–producing immune cells in intestinal defense against *C.r* was recently resolved^4^. While innate immune cells are effective at restraining bacterial colonization of the superficial epithelium early in infection, they are sequestered in isolated mucosal lymphoid tissues and primarily act at a distance^4, 45, 46^, becoming ineffective as infection progresses. On the other hand, after a delay required for the antigen-driven differentiation of Th17/Th22 effectors, these cells become the dominant producers of IL-22 following their migration to the infected distal colon. They are uniquely able to deliver sufficient IL-22 to IECs lining the colonic crypts as well as sustain IL-22 signaling into superficial IECs to maintain epithelial barrier function and to protect the crypts from pathogen encroachment^4^; in mice with deficiency of IL-22 targeted to T cells, pathogenic bacteria descend into the depths of crypts where they may damage epithelial stem cells that reside there. The ability of IL-22–producing T cells, but not innate immune cells, to induce much more robust, sustained STAT3 activation in IECs as infection progresses may reflect their localization in the LP immediately subjacent to both crypt and superficial IECs. How this T cell “help” for IECs is delivered is unclear, as are the epithelial cell populations affected. Moreover, it has been unclear why the geography of *C.r* infection is specifically limited to the mid-distal large intestine^12, 47, 48^.

Here, we describe discovery of a new absorptive IEC subpopulation that is regionally restricted to the mid-distal colon and is uniquely targeted by *C.r*, explaining *C.r*’s tropism for this tissue niche. We find that IL-22 signaling activates a distinct gene expression program in this new IEC subset compared to the conventional absorptive IEC subset that is dominant proximally in the large and small intestines. We further identify a critical role for MHCII expression by IECs to obtain IL-22 signaling from T cells required to protect colonic crypts from pathogen invasion, implicating an antigen-specific mechanism by which the intestinal epithelium elicits T cell help and stratifies the protective response. Thus, the host response to *C.r* infection reflects macro and micro geographical constraints on the pathogen and host tissue domains, and its success is contingent on MHC-restricted T cell help to IECs.

## Results

### Single-cell transcriptomics identify two major subsets of absorptive IECs with distinct developmental trajectories in response to *C.r* infection

Previous studies of the developmental and functional diversity of intestinal epithelial cell types have largely focused on the small intestine and have defined two major branches, absorptive and secretory^49–52^. In an effort to characterize large intestinal epithelial cell types as a prelude to exploring IEC responses induced by *C.r* infection, we performed single-cell (sc)RNA-seq analysis on IECs from the mid-distal colon, the region colonized by this pathogen (**Fig. 1**). Unexpectedly, we identified a bifurcation of the absorptive colonocyte lineage into two arms that we designated as distal colonocytes (DCC) and proximal colonocytes (PCC), for reasons to be discussed below (**Fig. 1a-d and Extended Data Figs. 1 and 3d**). These subsets derive from distinct early progenitors but their gene expression profiles identified them as members of the absorptive lineage, with both subsets expressing *Car4*, *Slc26a3*^53, 54^, and cell-surface mucins *Muc3* and *Muc13* that are main components of the protective glycocalyx layer that lies between goblet cell-derived secreted mucins (e.g., Muc2) and underlying colonocytes^55, 56^ (**Fig. 1a and Extended Data Fig. 1a,b and 3d**). However, mature DCCs uniquely expressed *Ly6g*, which encodes a surface marker for neutrophils that interacts with β2-integrins on leukocytes^57–59^, as well as acute phase proteins (e.g., *Saa1*, *Saa2*) that possess antimicrobial activity and can recruit and activate phagocytes^60–63^, and several solute carriers including *Slc20a1*, an inorganic sodium-phosphate cotransporter that can also regulate TNF-induced apoptosis^64, 65^ (**Fig. 1a,b and Extended Data Fig. 1d,e**). DCCs also expressed elevated levels of genes involved in metal ion storage (e.g., *Mt1*, *Fth1, Slc40a1*)^66–68^, suggesting that they may sequester metal ions to either reduce local metal toxicity and/or limit metal nutrient availability to commensal microbes. In contrast, developing and mature PCCs expressed *Fabp2*, an intracellular protein involved in lipid metabolism^69^, *Guca2a*, a polypeptide that activates guanylate cyclase receptors to control water transport in the intestines, and several glycosyltransferase genes that may be important for maintaining the mucosal barrier (**Fig. 1a-c and Extended Data Fig. 1c,f,g**). Mature DCC and PCC colonocytes also differed in expression of several SLC (solute carrier) transporters (e.g., DCCs express *Slc20a1* and *Slc40a1* whereas PCCs express *Slc6a14* and *Slc13a2*), which control the movement of various substances across their plasma membranes. Thus, *Ly6g*^+^ *Slc20a1*^+^ *Saa1*^+^ DCCs and *Fabp2*^+^ *Guca2a*^+^ *Reg3b*^+^ PCCs represent distinct subsets of the colonocyte absorptive lineage with gene expression profiles suggestive of functional specialization.

**Fig. 1:**
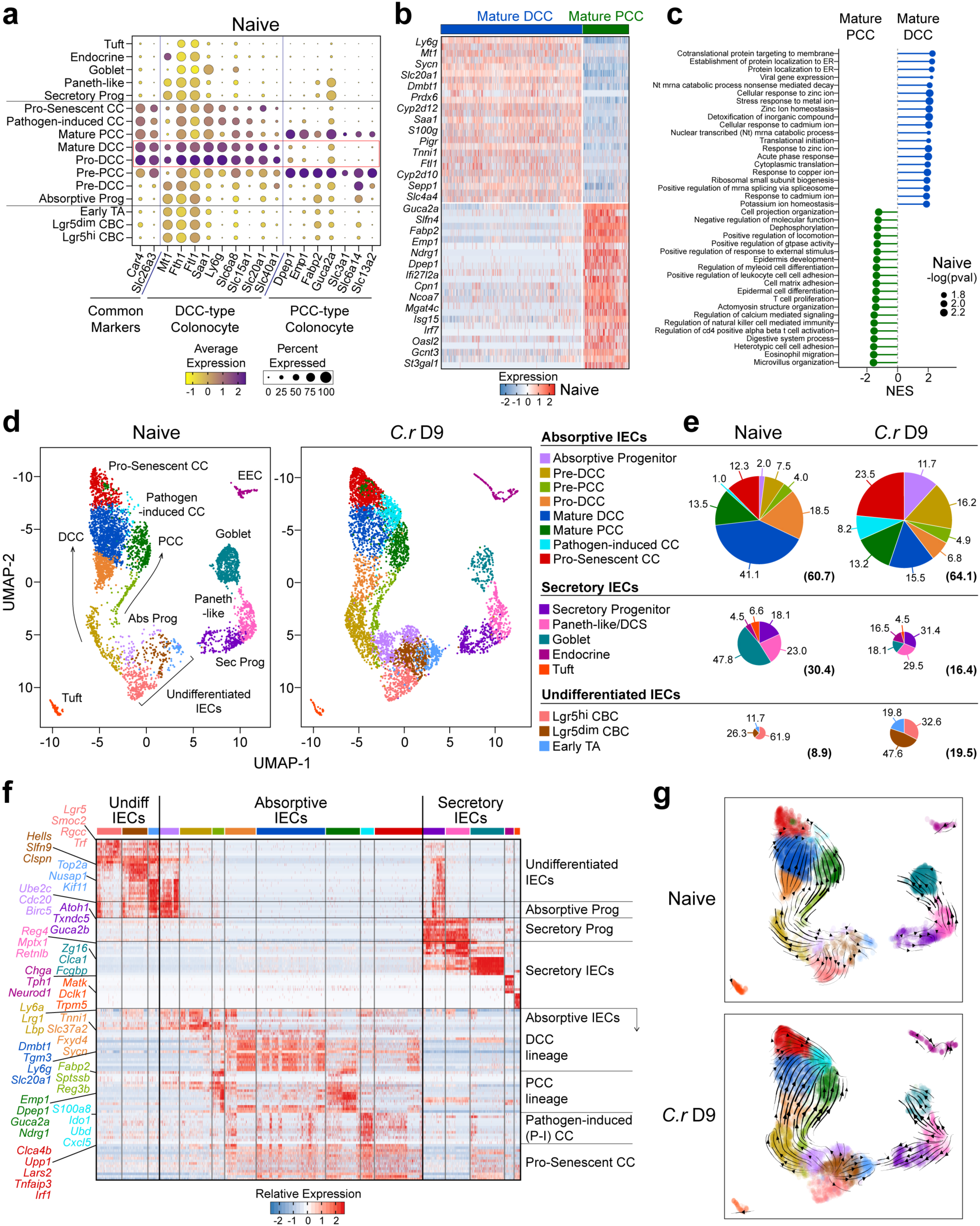
A distinct subset of colonocytes undergoes accelerated maturation in response to *C.r*. scRNA-seq was performed on colonic epithelial cells isolated from mid-distal colons of naïve and d9 *C.r*-infected BL/6 mice. **a**, Dot plot of DEGs and **b**, Heatmap of top 50 genes between DCC and PCC populations from naïve BL/6 mice. **c**, GSEA of GO-BP pathways comparing Mature DCC and Mature PCC from naïve BL/6 mice. **c**,. **d**, UMAPs of integrated biological replicates identified 8 unique cell clusters for absorptive IECs–including 2 major developmental branches (DCC and PCC), 5 clusters for secretory IECs and 3 clusters for undifferentiated IECs. **e**, Pie charts indicate percentages of cells within each major IEC subset. Numbers in parentheses indicate percentages of absorptive, secretory, and undifferentiated cells in each major pool of cells. **f**, Heatmap of top genes from combined naïve and infected mice defining each major IEC subset. **g**, scVelocity plots show transcriptional relationships between the major IEC subsets. Arrowheads denote directionality and lines represent kinetics of differentiation. 2 mice pooled per sample, *n*=2 biological replicates per group. IEC=intestinal epithelial cell; CC=colonocyte; PCC=middle colonocyte; DCC=distal colonocyte; DCS=deep crypt secretory cell; TA=transit-amplifying cell; CBC=crypt base columnar cell; Abs= absorptive IEC; Sec=secretory IEC; Undiff=undifferentiated cell; Prog=progenitor; NES=normalized enrichment score.

Colonic hyperplasia—elongation of the crypts associated with expansion of undifferentiated crypt base columnar (CBC) stem cells, transit-amplifying (TA) cells and progenitor IECs—is a hallmark of *C.r* infection^12, 13, 48, 53^ It is thought to be important as a means to protect the stem cells at the base of the crypts by both distancing these cells from the pathogen and accelerating removal of *C.r*-laden IECs beyond the mouth of the crypts^52, 70–74^ (**Extended Data Fig. 2a,b**). To characterize the hyperplastic response of IEC subsets in the region colonized by *C.r* infection, we extended our scRNA-seq analysis to compare IECs isolated from the mid-distal colon of naïve versus *C.r*-infected mice (**Fig. 1d-g**). Strikingly, DCCs and PCCs differed in their response to *C.r* infection (**Fig. 1d-f and Extended Data Fig. 2c**). Progenitors of DCCs underwent considerable expansion whereas PCCs showed modest changes in frequency, abundance or developmental programming (**Fig. 1d-f and Extended Data Fig. 2c**). Differentiated DCCs (pro-DCC and mature DCC) were reduced with a concomitant increase in a population of “pathogen-induced colonocytes” (P-ICC) that express IL-22–inducible and IFNγ–inducible host defense genes (e.g., *S100a8*, *Cxcl5* and *Ido1*)^1, 17, 18, 75, 76^ and colonocytes with a gene profile resembling a pro-senescent state (e.g., *Upp1*, *Lars2*, *Tnfaip3*)^77–81^ (**Fig. 1d-f and Extended Data Table 1**). scRNA velocity analysis indicated that in response to *C.r* infection, DCC lineage cells underwent rapid maturation and an altered cell fate to become “hyperactive” P-ICCs that contribute to host defense (**Fig. 1g**).

Undifferentiated IECs (Lgr5^+^ CBCs^82^ and early TA cells) expanded, while goblet cells were reduced in response to *C.r* infection (**Fig. 1d-e**). This is consistent with previous reports implicating IFNγ-producing CD4 T cells in control of stem cell proliferation and goblet cell reprogramming and hypoplasia during *C.r* infection^83, 84^. In addition, some colonic *Reg4*^+^ *Mptx1*^+^ Paneth-like cells (also called deep crypt secretory cells)^85, 86^ dedifferentiated during *C.r* infection, paralleling plasticity in small intestinal Paneth cells that transition into CBC-like stem cells in response to inflammation and injury^87, 88^. These findings identify unanticipated heterogeneity within the colonocyte absorptive lineage and implicate the newly identified DCC subset as the dominant responder to *C.r* infection. They further indicate that a developmental shift in colonic Paneth-like cells may contribute to the reduction in goblet cells (hypoplasia), and expansion of DCC progenitors during *C.r* infection. Collectively, then, these results establish that colonic hyperplasia in response to *C.r* infection predominantly comprises rapid expansion of DCC lineage cells to generate P-ICCs with heightened expression of cytokine-induced host defense molecules.

### DCCs are restricted to distal colon and are the principal target of *C.r.* infection

The foregoing studies identified differential responses of DCCs and PCCs to *C.r* but did not identify the basis. We considered that this might reflect geographic segregation of these two subsets, differential interactions with *C.r*, or both. To explore the first possibility, we evaluated the distribution of DCCs and PCCs along the intestinal tracts of naïve mice, employing subset-specific markers that had been identified from our scRNA-seq studies; Ly6g was used as a DCC-specific marker and Fabp2 as a PCC-specific marker (**Fig. 2a,b and Extended Data Fig. 1d-g**). Remarkably, we found that Ly6g^+^ cells (DCCs) were found exclusively in the distal colon and were interspersed with Fabp2^+^ cells (PCCs) in the middle colon, while Fabp2^+^ cells dominated proximally in colon and in terminal ileum of small intestine (**Fig. 2c-e**). Hence, the designations “distal” and “proximal” colonocytes.

**Fig. 2:**
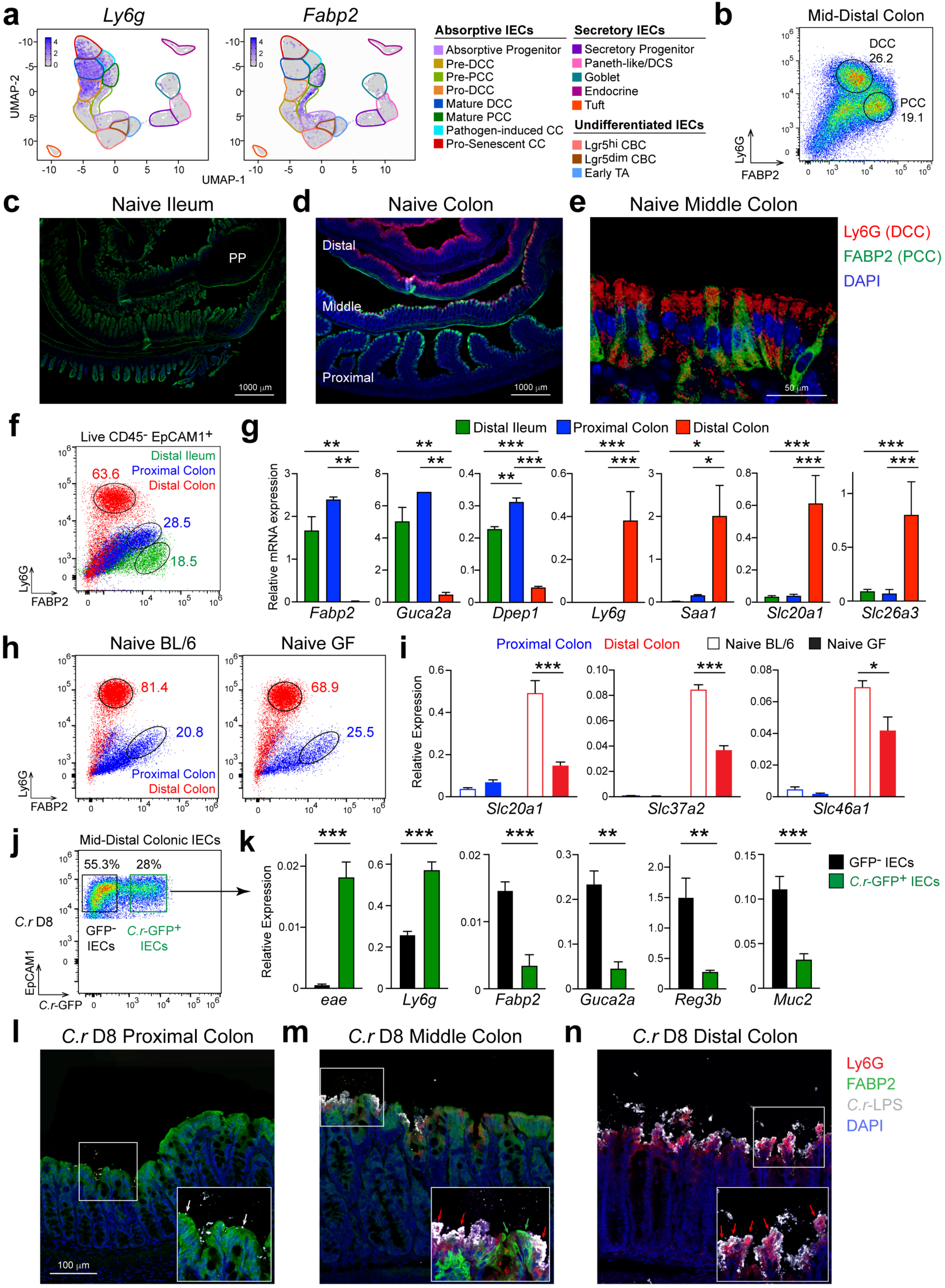
*Citrobacter rodentium* predominantly attaches to DCCs. scRNA-seq was performed on colonic epithelial cells isolated from mid-distal colons of naïve and d9 *C.r*-infected BL/6 mice. **a**, UMAP of *Ly6g* and *Fabp2* genes. 2 mice pooled per sample, 2 biological replicates per group for scRNA-seq. **b**, IECs from mid-distal colon of naïve BL/6 mice were stained for Ly6G, Fabp2, EpCAM1, CD45 and L/D dye, and analyzed by flow cytometry. 3 mice per experiment and *n*=2 independent experiments. **c-e**, Tissue from distal ileum; scale bar, 1000 μm (**c**), colon; scale bar, 1000 μm (**d**) and middle colon region; scale bar, 50 μm (**e**) from naïve BL/6 mice were stained with Ly6G (red; DCC), Fabp2 (green; PCC) and DAPI (blue). 3-4 mice per region and *n*=2 independent experiments. **f-g**, IECs from distal ileum (green), proximal colon (blue) and distal colon (red) of naïve BL/6 mice were stained with Ly6G, Fabp2, EpCAM1, CD45 and L/D dye, and analyzed by flow cytometry (**f**) or sorted on EpCAM1^+^ CD45^-^ L/D dye^-^ IECs and *Fabp2*, *Reg3b*, *Ly6g*, *Slc20a1* and *Slc26a3* mRNA expression analyzed by RT-PCR (**g**). 2-3 mice pooled per sample, *n*=2-3 biological replicates per group and *n*=2 independent experiments. One-way ANOVA; *p≤0.05, **p≤0.01 and ***p≤0.001 comparing different intestinal regions. **h-i**, IECs from proximal colon (blue) and distal colon (red) of naïve BL/6 and germ-free (GF) mice were stained with Ly6G, Fabp2, EpCAM1, CD45 and L/D dye, and analyzed by flow cytometry (**h**) or sorted on EpCAM1^+^ CD45^-^ L/D dye^-^ IECs and *Slc20a1*, *Slc37a2* and *Slc46a1* mRNA expression analyzed by RT-PCR (**i**). 3-4 mice per group and *n*=2 independent experiments. Two-way ANOVA; *p≤0.05 and ***p≤0.001 comparing BL/6 and GF mice. Data are represented as mean ± SEM. **j**, Mid-distal colon IECs from d8 *C.r*-GFP-infected mice were sorted on either EpCAM1^+^ CD45^-^ GFP^-^ (black) or EpCAM1^+^ CD45^-^ *C.r*-GFP^+^ cells (green). **k**, Sorted *C.r*-GFP^-^ and *C.r*-GFP^+^ IECs were analyzed for *eae*, *Ly6g, Fabp2, Guca2a, Reg3b* and *Muc2* mRNA expression by RT-PCR. 2-3 mice pooled per sample, *n*=2-3 biological replicates per group and *n*=2 independent experiments. Student’s t-test; **p≤0.01 and ***p≤0.001 comparing *C.r*-GFP^-^ IECs and *C.r*-GFP^+^ IECs. **l-n**, Proximal (**l**), middle (**m**) and distal (**n**) colon tissue from d8 *C.r*-infected mice was stained for Ly6G (red) or Fabp2 (green), *C.r*-LPS (white) and DAPI (blue). White arrows depict few *C.r* attached in proximal colon, green arrows depict untouched Fabp2^+^ PCCs, and red arrows depict *C.r*-laden Ly6G^+^ DCCs. Scale bar, 100 μm. 3-4 mice per experiment and *n*=2 independent experiments. IEC=intestinal epithelial cell; CC=colonocyte; PCC=middle colonocyte; DCC=distal colonocyte DCS=deep crypt secretory cell; TA= transit-amplifying cell; CBC=crypt base columnar cell.

To confirm these findings, IECs from distal ileum, proximal colon or distal colon were isolated by flow cytometric sorting and their gene expression profiled by RT-PCR based on scRNA-seq analysis of mature DCCs and PCCs. Epithelial cells isolated from the ileum and proximal colon expressed *Fabp2*, *Guca2a* and *Dpep1*—signature PCC-type genes; in contrast, cells from the distal colon expressed *Ly6g*, *Saa1* and *Slc20a1*—signature DCC genes (**Fig. 2f,g and Extended Data Figs. 1e-g and 2c**). Interestingly, *Slc26a3*, a chloride/bicarbonate transporter associated with congenital chloride diarrheal disease^89, 90^ appears to be a specific marker of mid-distal colonocytes and is not expressed in the proximal colon or terminal ileum (**Fig. 2g and Extended Data Fig. 1a,b**).

To determine if the development of either or both of these IEC subsets might be driven by interactions with the commensal microbiota, we assessed their relative abundance in naive conventionally housed specific pathogen-free (SPF) and germ-free (GF) mice (**Fig. 2h**). There was no significant difference in either population, indicating that the development of both is largely independent of signals from the commensal flora. However, several DCC-specific genes were upregulated in DCCs contingent on the commensal microbiota, including SLC family members *Slc20a1*, *Slc37a2* and *Slc46a1* (**Fig. 2i**). Collectively, these data indicate that DCCs represent a newly identified subset of absorptive IECs that are regionally restricted to mid-distal colon. Although their development is independent of signals from the commensal microbiota, DCCs appear to be constitutively responsive to commensals and may sequester metal nutrients to maintain the barrier in this region.

In view of the striking regionalization of DCCs to the same area of the colon colonized by *C.r*^3, 4, 12, 13, 47^, and the preferential infection-induced expansion of DCC progenitors and reciprocal reduction in mature DCCs (**Fig. 1**), we speculated that mature DCCs may be targets of *C.r* colonization. To test this directly, we took advantage of the tight association of *C.r* to infected IECs and a GFP-expressing *C.r* variant^91, 92^ to sort isolated uninfected and infected IECs from mid-distal colons of WT mice and screen for DCC- and PCC-specific genes (**Fig. 2j,k and Extended Data Fig. 3a,b**). As a control and internal validation, the *C.r*-specific gene *eae,* which encodes intimin, a molecule crucial for *C.r* attachment to IECs^10, 93^, was strictly restricted to *C.r*-GFP-laden IECs. The *Ly6g* gene, expression of which is specific for DCCs, was significantly enriched in infected *C.r*-GFP^+^ IECs but was also found in uninfected *C.r*-GFP^−^ IECs, establishing that *C.r* binds directly to DCCs but also suggesting that there is non-uniform coverage of DCCs. The latter likely reflects lack of *C.r* attachment to some mature DCCs and pro-DCCs that express Ly6g but are sequestered within crypts, whereas *C.r*-GFP^+^ IECs are likely infected mature DCCs and pro-senescent DCCs that are fated to be shed and removed from the colon. In contrast, genes that define PCCs (e.g., *Fabp2*, *Guca2a*, *Reg3b*) or goblet cells (e.g., *Muc2*) were found predominantly in the uninfected subpopulation of IECs, indicating that most of these IECs are not bound by *C.r*.

To extend these findings, we examined *C.r* attachment to either Ly6g^+^ DCCs or Fabp2^+^ PCCs in the colons of *C.r*-GFP-infected WT mice *in situ* (**Fig. 2l-n**). There was minimal *C.r* colonization of the proximal colon. In the middle colon, which is populated by both DCCs and PCCs, *C.r* attached to Ly6g^+^ DCCs with few Fabp2^+^ PCCs targeted by *C.r*. In the distal colon, where the absorptive epithelium is comprised almost exclusively of DCCs, the majority of superficial Ly6g^+^ DCCs were bound by *C.r,* demonstrating that *C.r* preferentially colonizes DCCs found in the mid-distal colon. Thus, *C.r* shows strong tropism for mature DCCs, explaining the restriction of colonization by this pathogen to the distal colon.

### Proximal and distal colonocytes respond differentially to T cell-derived IL-22

Based on the differential targeting of subsets of absorptive colonocytes by *C.r*, we speculated that there might be stratification of host defense responses between DCCs and PCCs. Comparison of the gene expression profiles of DCC and PCC cells from mid-distal colons of WT mice by scRNA-seq indicated that this was indeed the case. While both DCCs and PCCs downregulated a considerable number of genes during *C.r* infection (e.g., *Car4*, *Slc26a3*, and several differentially expressed solute carrier genes)—indicating that both colonocyte subsets sensed the pathogen and altered their response^53, 54, 94, 95^, a preponderance of genes that we had previously reported as contingent on T cell-derived IL-22^4^ were specifically upregulated in DCCs (**Fig. 3a,b and Extended Data Fig. 5**). Notably, the *S100a* family of antimicrobial peptides (AMPs; e.g., *S100a8*) and neutrophil-recruiting chemokines (e.g., *Cxcl2* and *Cxcl5*) were strongly upregulated on IECs of the DCC lineage (pro-DCC, mature DCC and P-ICC) in infected control compared to naïve mice.

**Fig. 3:**
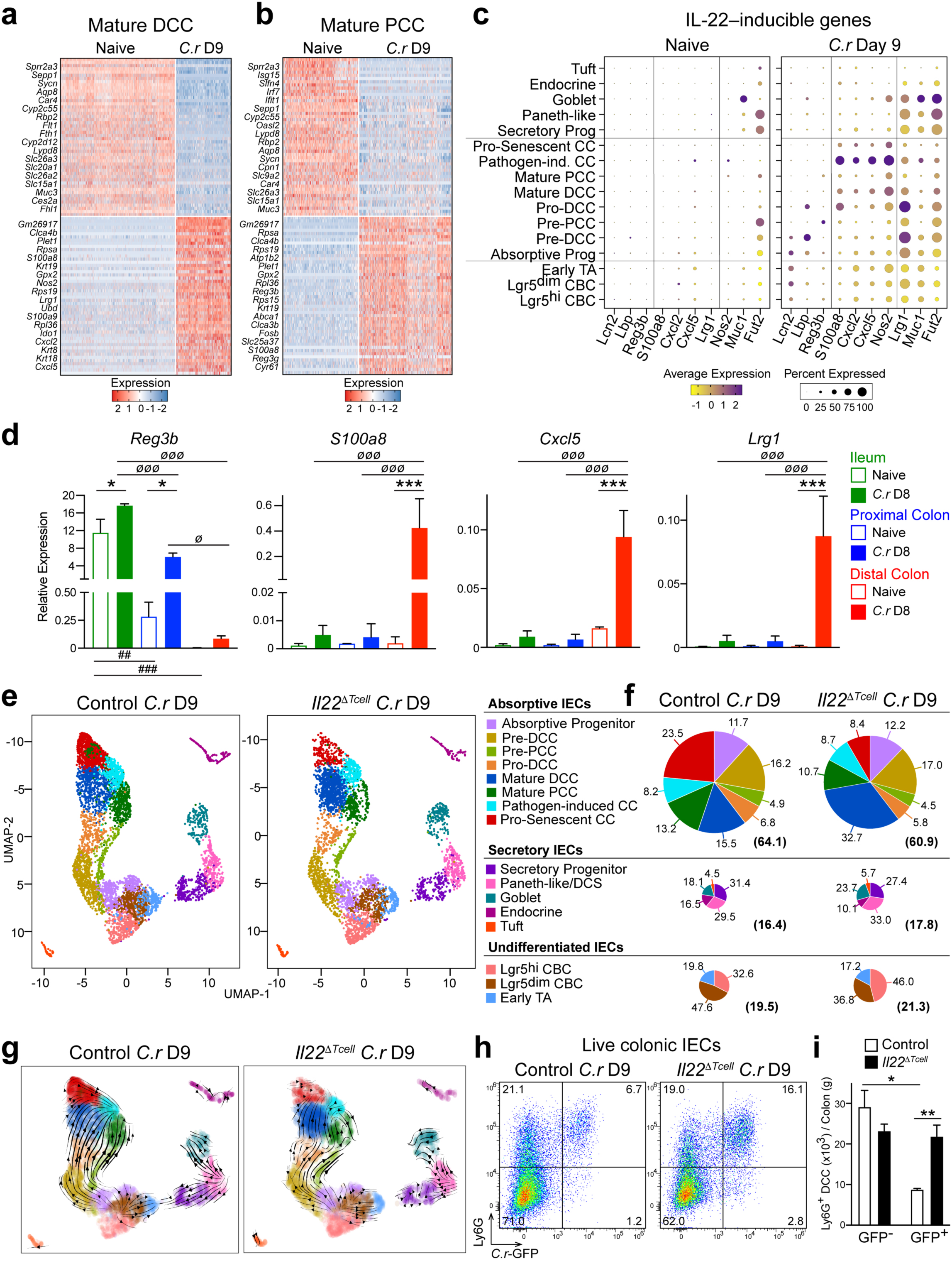
IL-22^+^ T cells accelerates removal of *C.r*-laden mature DCCs. scRNA-seq was performed on colonic epithelial cells isolated from mid-distal colons of naïve BL/6 and d9 *C.r*-infected *Il22*^hCD4^ (control) and *Il22*^ΔTcell^ cKO mice. **a-b**, Heatmap of top 50 genes expressed by Mature DCC (**a**) and Mature PCC (**b**) comparing naïve and infected mice. **c**, Dot plot of IL-22-inducible genes expressed by colonic IECs from naïve and infected mice. 2 mice pooled per sample, 2 biological replicates per group for scRNA-seq. **d**, IECs from naïve (open symbol) and d8 *C.r*-infected mice (closed symbol) were sorted from distal ileum (green), proximal colon (blue) and distal colon (red) and *Reg3b*, *S100a8*, *Cxcl5* and *Lrg1* mRNA expression analyzed by RT-PCR. One-way ANOVA with Tukey’s multiple comparison test; *p≤0.05, and ***p≤0.001 comparing naïve and infected mice; ^ø^p≤0.05 and ^øøø^p≤0.001 comparing infected samples from different tissue regions; and ^##^p≤0.01, and ^###^p≤0.001 comparing naïve samples from different tissue regions. 2-3 mice pooled per sample and *n*=2 independent experiments. Data are represented as mean ± SEM. **e**, UMAPs of integrated biological replicates from d9 *C.r*-infected control and *Il22*^ΔTcell^ cKO mice. **f**, Pie charts indicate percentages of cells within each major IEC subset. Numbers in parentheses indicate percentages of absorptive, secretory, and undifferentiated cells in each major pool of cells. **g**, scVelocity cell cluster plots showing transcriptional relationships between the major IEC subsets. Arrowheads denote directionality and lines represent kinetics of differentiation. 2-3 mice pooled per sample, *n*=2 biological replicates per group for scRNA-seq. **h**, IECs isolated from colons of d9 *C.r*-GFP-infected *Il22*^hCD4^ and *Il22*^ΔTcell^ mice were stained for Ly6G, CD45, L/D dye and EpCAM1 and analyzed by flow cytometry. **i**, Number of *C.r*-GFP^-^ Ly6G^+^ IECs and *C.r*-GFP^+^ Ly6G^+^ IECs from d9 *C.r*-GFP-infected *Il22*^hCD4^ (white) and *Il22*^ΔTcell^ (black) mice. Student’s t-test; *p≤0.05 and **p≤0.01 comparing *Il22*^hCD4^ and *Il22*^ΔTcell^ mice. 3-4 mice per group and *n*=2 independent experiments. Data are represented as mean ± SEM. IEC=intestinal epithelial cell; CC=colonocyte; PCC=middle colonocyte; DCC=distal colonocyte; DCS=deep crypt secretory cell; TA=transit-amplifying cell; CBC=crypt base columnar cell.

A specific contribution of the DCC lineage to host defense appeared to be the unique production of neutrophil-recruiting chemokines required for *C.r* eradication^96, 97^ (**Fig. 3a-c**). Particularly striking was the IL-22–induced enrichment of these chemokine transcripts in the P-ICC subpopulation, in which the greatest frequency and amplitude of expression of these genes was found (**Fig. 3c and Extended Data Figs. 3e and 4**). In addition, pro-DCC, mature DCCs and DCC lineage-derived P-ICCs from infected mice specifically upregulated *Nos2*, which encodes the inducible enzyme nitric oxide (NO) synthase important for production of NO, a broad-spectrum antimicrobial molecule that causes nitrosative and oxidative cell damage^98, 99^. Furthermore, immature subsets of DCCs upregulated IL-22–inducible *Lcn2*, *Lbp* and *Fut2,* indicating that colonocytes in the mid-lower crypts are also capable of sensing and responding to microbial antigens during *C.r* infection. In contrast, upregulation of the IL-22-inducible *Reg3* lectin family of AMPs by *C.r* infection was PCC-specific. Because *C.r*-infected *Reg3b*^−/−^ mice have enhanced bacterial load^100, 101^ this suggests that PCCs may contribute to host defense by reducing luminal bacteria that can attach to DCCs, although other mechanisms, including modulation of the commensal microbiota, cannot be excluded.

In accord with these findings, IL-22–induced transcripts of IECs isolated from either distal colon, proximal colon, or distal ileum of naïve and *C.r*-infected WT mice correlated well with the distribution of DCCs and PCCs across these intestinal segments (**Fig. 3d and Extended Data Fig. 3c**). Consistent with published reports, we found that *Reg3b* was predominantly expressed in the small intestine with low levels of *Reg3b* expression in the proximal colon at steady state^101^. At the peak of *C.r* infection (d8-9), *Reg3b* was upregulated on IECs from the distal ileum and proximal colon but not the distal colon. In contrast, other IL-22–inducible genes such as *S100a8*, *Cxcl5* and *Lrg1* were predominantly upregulated in the distal colon but not ileum or proximal colon. Together these data establish that upregulation of *S100a*8/9 AMPs and neutrophil-attractant chemokines reflect the direct targeting by *C.r* of DCCs that populate the distal colon, whereas PCCs upregulate Reg3 family AMP transcripts that may augment pathogen clearance in the lumen. Whether the IL-22–responsive PCCs are further regionalized in proximity to the DCCs at the intersection of middle and distal colon is not yet known, but the relatively minor fraction of PCCs that strongly express these transcripts is suggestive that this is plausible. In any event, DCCs and PCCs appeared to be programmed to work in concert to coordinate host defense against *C.r*^96, 102–104^, albeit via distinct biological activities downstream of IL-22.

scRNA-seq analysis of mid-distal IECs from *C.r*-infected *Il22*^fl/fl^ (control) and *Il22*^ΔTcell^ (T cell-specific deficiency of IL-22) confirmed this (**Fig 3e,f and Extended Data Figs. 3d,e and 5**); the induction of S100 family AMPs and neutrophil-attractant chemokines or Reg3 family AMPs was DCC- or PCC-specific, respectively, and contingent on actions of T cell-derived IL-22 (**Extended Data Figs. 3d,e, 4 and 5**). In contrast, and similar to the expansion of stem cells and reprogramming of goblet cells, which was comparable between *C.r*-infected control and *Il22*^ΔTcell^ mice and consistent with their regulation by IFNγ^83, 84^, not IL-22, the expansion of absorptive progenitors and immature pre-DCCs was not controlled by T cell-derived IL-22. Moreover, scRNA velocity analysis indicated that the transition from immature to mature colonocyte subsets during *C.r* infection in control mice progressed in a constant acceleration pattern (**Fig. 3g**). However, we found a critical role for T cell-derived IL-22 to program DCCs to accelerate their transition to pro-senescence during *C.r* infection (**Fig. 3e-g**), and the migration of maturing DCC subsets in *C.r*-infected *Il22*^ΔTcell^ mice was irregular (**Fig. 3g**), suggesting discontinuous terminal differentiation. The aberrant transition of DCCs to pro-senescent colonocytes in *C.r*-infected *Il22*^ΔTcell^ mice compared to controls suggested that IL-22–producing T cells act to promote the more rapid removal of *C.r*-laden colonocytes.

This proved to be the case, as we found a significant increase in live *C.r*-GFP^+^ DCCs in the colons of *C.r-*GFP-infected *Il22*^ΔTcell^ mice compared to controls (**Fig. 3h,i**). In addition, compared to *C.r*-GFP-infected *Il22*^ΔTcell^ mice, control mice had significantly higher numbers of GFP^−^ DCCs compared to GFP^+^ DCCs, further implicating IL-22 produced by T cells in promoting replacement of *C.r*-laden DCCs with newly generated DCC lineage cells. In this regard, we found it notable that T cell-derived IL-22 augmented *Lrg1* expression at multiple stages of DCC development. *Lrg1*, which encodes for a leucine-rich α-2-glycoprotein involved in epithelial cell migration and wound healing^105, 106^, was upregulated by most IECs during *C.r* infection but its greatest expression was found in immature DCCs (pre-DCC and pro-DCC), suggesting that *Lrg1* may play a role in the movement of distal colonocytes up the crypts during *C.r* infection (**Fig. 3c,d and Extended Data Figs. 2c, 3e and 5**). Together these findings indicate that the delivery of IL-22 to DCCs by CD4 T cells is required to both activate these cells for enhanced anti-bacterial defense and accelerate the removal of *C.r*-laden colonocytes and their replacement at the luminal surface to expedite pathogen clearance, the latter representing a novel mechanism by which IL-22 signaling into DCCs may counter *C.r*-mediated effector mechanisms to promote the retention of infected DCCs to which they anchor^107–109^.

### Epithelial expression of MHCII is required to elicit protective help from CD4 T cells

The injection of bacterial effector proteins into host IECs is required for the attachment of *C.r* to superficial DCCs that line the surface of the distal colon. This also represents a potential vulnerability for *C.r*, as these effector proteins may be sensed by intracellular pattern-recognition receptors and provide a source of antigenic peptides that can be recognized by T cells. Although antigenic priming of naive CD4 T cells in response to *C.r* is thought to occur in lymphoid tissues by type 2 conventional dendritic cells (cDC2s)^27, 110^, it is unclear whether *C.r* antigens can be presented by IECs to directly recruit T cell help. In a previous study, we found that IECs begin to upregulate MHCII and other components of the class II antigen processing system coincident with the influx of effector CD4 T cells into the infected mid-distal colon at the peak of *C.r* colonization (d7-9), and the majority of IECs express MHCII later in infection (d12-20)^4, 54^, concurrent with sustained STAT3 activation in IECs driven by T cell-derived IL-22 signaling^4, 54^. The coordination of timing of IEC upregulation of the class II antigen processing and presentation system with the recruitment of robust, T cell-dependent IL-22 signaling into IECs led us to posit a role for direct, antigen-dependent release of IL-22 by *C.r-*specific Th17/Th22 cells.

To test this, we explored both T cell and IEC responses to *C.r* infection in mice with specific deficiency of MHCII targeted to epithelial cells (*H2-Ab1*^Villin^). We found that 90% of Ly6g^+^ DCC and Fabp2^+^ PCC cells from mid-distal colon upregulate MHCII compared to the 20% of Fabp2^+^ proximal colonocytes and distal ileal enterocytes on d14 of *C.r* infection (**Fig. 4a,b**), indicating that IFNγ, which is required for induction of the MHCII system^111–113^, was elevated in both the middle and distal colon compared to the proximal colon and SI during the adaptive phase of *C.r* infection. On infection d9, when MHCII expression is upregulated on a small fraction of IECs^4^, there was no difference in the number of Th17 and Th22 cells in the colons of infected *H2-Ab1*^Villin^ compared to *H2-Ab1*^fl/fl^ control mice, consistent with a dispensable role for epithelial MHCII in the differentiation and recruitment of effector T cells at this early phase of the adaptive response to infection (**Fig. 4c,d**). In contrast, by infection d14, when the majority of mid-distal colonocytes expressed MHCII in control mice, there was >50% reduction in the numbers of Th17 and Th22 cells in the colons of infected *H2-Ab1*^Villin^ mice compared to controls. This correlated with comparable *C.r* burdens in fecal contents and liver on d9 but significantly heightened burdens on d14 post-infection of infected *H2-Ab1*^Villin^ mice compared to controls (**Fig. 4e**). Moreover, compared to controls mice deficient for MHCII on IECs (*H2-Ab1*^Villin^) showed a striking loss of STAT3 activation in both crypt-lining and superficial IECs during the late phase of *C.r* infection (**Fig. 4f**). This was accompanied by invasion of the colonic crypts by *C.r* in *H2-Ab1*^Villin^ mice but not control mice (**Fig. 4g**), phenocopying the defect in mice with T cell-specific IL-22 deficiency to activate STAT3 and defend the colonic crypts against bacterial invasion^4^. Collectively, these findings establish a requirement for MHCII expression by IECs of the distal colon to retain Th17 and Th22 cells and elicit IL-22-dependent help from these cells that is indispensable for host defense against *C.r*.

**Fig. 4:**
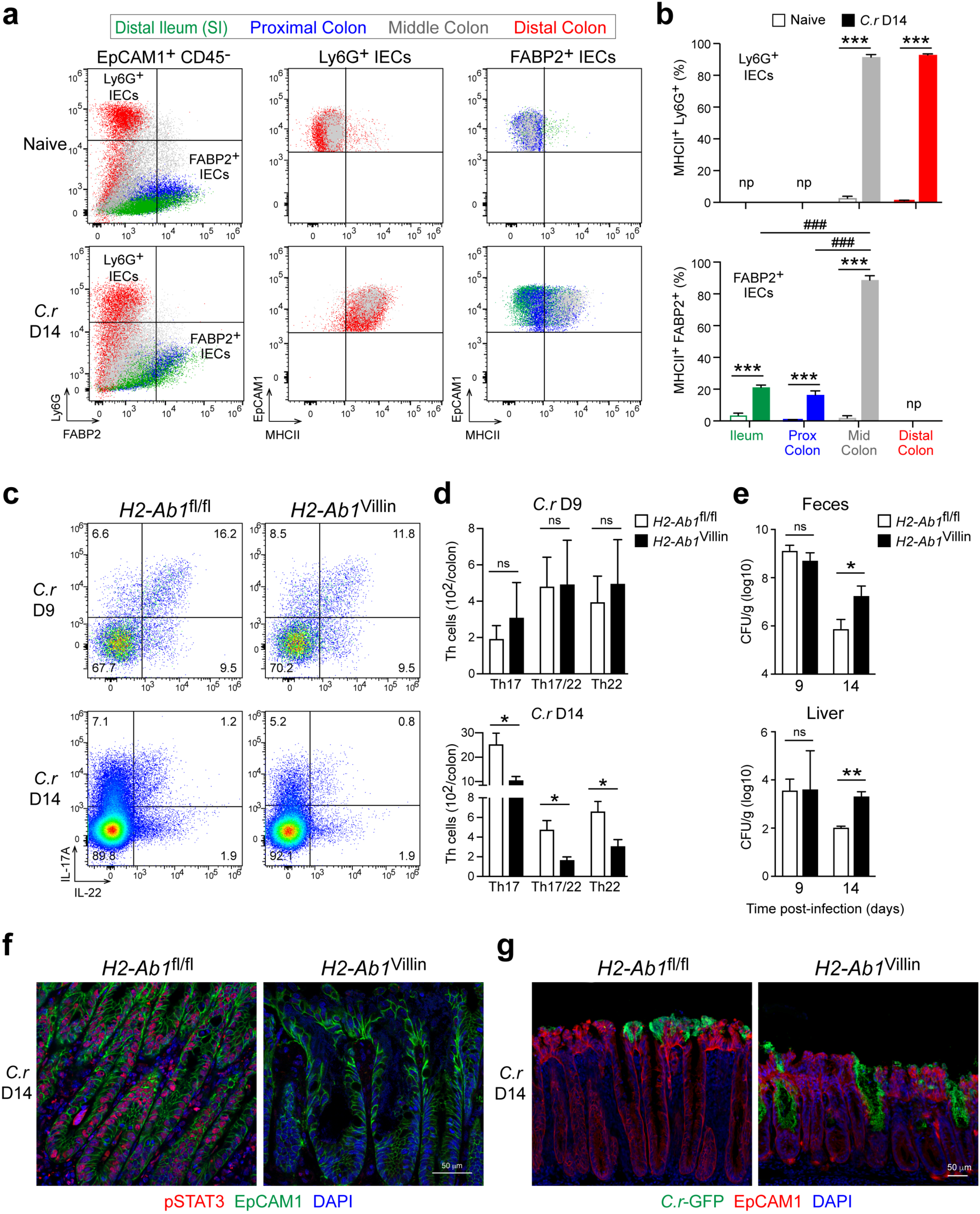
Epithelial MHCII maintains colonic Th cells for prolonged STAT3 activation and crypt protection. **a**, IECs from distal ileum (green), proximal colon (blue), middle colon (grey) and distal colon (red) of naïve and d14 *C.r*-infected BL/6 mice was stained for EpCAM1, CD45, LD dye, Ly6G, Fabp2 and MHCII and analyzed by flow cytometry. 3 mice pooled per region, *n*=2 independent experiments. **b**, Percent of MHCII^+^ Ly6G^+^ DCCs and MHCII^+^ Fabp2^+^ IECs from naïve and d14 C.r-infected BL/6 mice. Two-way ANOVA; *p≤0.05, **p≤0.01 and ***p≤0.001 comparing naïve versus infected mice. **c**, Colon cells from d9 and d14 *C.r*-infected *H2-Ab1*^fl/fl^ floxed and *H2-Ab1*^Villin^ cKO mice were stimulated for 4hrs with PMA/Ion and Brefeldin, stained for surface CD4, TCRβ and L/D dye, and then stained for intracellular IL-17 and IL-22 and analyzed by flow cytometry. **d**, Number of colonic Th17, Th17/22 and Th22 cells from d9 and d14 *C.r*-infected *H2-Ab1*^fl/fl^ (white) and *H2-Ab1*^Villin^ (black) mice. 3-4 mice per group, *n*=2 independent experiments. **e**, Log_10_ CFU of d9 and d14 *C.r*-GFP in feces and liver from infected *H2-Ab1*^fl/fl^ (white) and *H2-Ab1*^Villin^ (black) mice. 4-5 mice per group, *n*=2 independent experiments. Student’s t-test; *p≤0.05 and **p≤0.01 comparing *H2-Ab1*^fl/fl^ and *H2-Ab1*^Villin^ mice. **f-g**, Colon tissue from d14 *C.r*-GFP-infected *H2-Ab1*^fl/fl^ and *H2-Ab1*^Villin^ mice was stained for (**f**) pSTAT3 (red), EpCAM1 (green) and DAPI (blue) or (**g**) *C.r*-GFP (green), EpCAM1 (red) and DAPI (blue). 3-5 mice per group and *n*=2 independent experiments. Scale bar, 50 μm. Data are represented as mean ± SEM. ns=not significant. np=not present.

## Discussion

A fundamental property of effector CD4 T cells is their provision of antigen-dependent “help” to cells that express cognate antigen. Here, we identify a new subset of absorptive colonocytes in the mid-distal colon that are the cellular host of the murine enteropathogen, *Citrobacter rodentium*, and show that these cells obtain IL-22 from effectors of the Th17 pathway via MHCII-restricted interactions to protect the epithelium and crypts from pathogen invasion. Our findings address the longstanding conundrum of why *C.r* colonization is regionally restricted^47^—*C.r* attachment is primarily limited to the middle and distal colon due to its tropism for DCCs. Our findings also reveal a basis for the indispensable role of T cells in protection of the intestinal barrier as infection progresses and IL-22 derived from innate immune cells becomes less effective: T cells acting locally through MHCII-restricted interactions with the intestinal epithelium are required to sustain high amplitude IL-22 signaling to IECs as they undergo developmental shifts in response to infection. Finally, our findings provide a basis for a requirement of coordinated signaling of IL-22 and IFNγ into IECs under pathogen threat. Because IFNγ is required to induce the MHCII-restricted antigen processing pathway in IECs, it is prerequisite for the protective actions of IL-22—or other signals that may be delivered directly to IECs by T cells; in essence, IFNγ signaling “licenses” IECs for recruitment of enhanced IL-22 signaling from Th17/Th22 cells, which is essential for barrier protection.

In contrast to *Citrobacter*, which targets DCCs, ileal enterocytes are the infectious niche for EPEC in humans^114^. The specific host molecule(s) required for initial attachment of *C.r* or enteropathogenic *E.coli* to IECs is unknown. Although firm attachment to IECs by both pathogens relies on intimin, an outer membrane adhesion molecule that mediates bacterial attachment to IECs via the translocated intimin receptor (Tir), it is thought that fimbriae and pili mediate initial attachment to IECs via exposed D-mannose residues on glycoproteins^115, 116^. However, intimin may also contribute to bacterial tropism. Most EPEC strains express type α intimin, whereas *C.r* uses intimin-β^93,117^. Thus, in addition to species-specific differences in expression of fimbriae and pili, sequence differences between bacterial intimin subtypes may contribute to host and tissue specificity. Our discovery of a new subset of absorptive colonocytes that are cellular targets for *C.r* colonization should facilitate comparative studies to define the basis for species- and cell-specific tropism of A/E enteropathogens.

The identification of colonocyte subset-specific programming by IL-22 has implications for host anti-bacterial defense strategies. Notably, IL-22–dependent regulation of the Reg3 family of defensins originally implicated in antibacterial defense against *C.r*^1^ is restricted to PCCs, which are not targeted by *C.r*. This suggests indirect rather than direct actions of this family of AMPs in host protection. Nevertheless, Reg3 AMPs expressed by PCCs would appear to contribute to resistance to *C.r* infection and the clearance of *C.r*-laden DCCs^100, 118^, through mechanisms yet to be defined. In contrast, DCCs utilize IL-22 delivered by T cells to upregulate genes such as *lipocalin 2* (*Lcn2*)^119, 120^ that bind to bacterial siderophores which sequester iron from the host, and the antimicrobial peptide S100a8/9 (calprotectin)^121, 122^, a calcium/zinc-binding metal chelator that prevents bacterial uptake of metal ions during *C.r* infection. In addition, IL-22–stimulated mature DCCs upregulate Cxcl2 and Cxcl5 chemokines in response to infection to recruit neutrophils that themselves produce Lcn2 and S100a8/9^123, 124^, thereby amplifying local antimicrobial activity. In this regard, it is notable that several genes involved in metal ion homeostasis (e.g., *Ftl1*, *Slc40a1*) are expressed by DCC lineage cells at steady state and are downregulated during *C.r* infection. The acquisition of metal ions by *E.coli* and *C.r* is a double-edged sword—while it benefits bacterial growth, dysregulation of iron availability can also generate toxic reactive oxygen species that damages the epithelial cells they need to colonize. Insights into how these mechanisms operate to favor host or pathogen will be facilitated by findings herein.

*Citrobacter* and other A/E enteropathogens must attach to survive. In addition to induction of AMPs and neutrophil-attractant chemokines, a novel mechanism of host resistance mediated by T cell-delivered IL-22 was found to be programming of DCCs for accelerated clearance from the epithelium, presumably to counter actions of bacterial effector molecules injected into *C.r*-bound DCCs that act to thwart programmed senescence of IECs to prolong survival of *C.r*’s cellular host^107, 125, 126^. The retention of pro-senescent DCCs in the absence of T cell-derived IL-22 is particularly striking given that decreased shedding of colonized DCC cells increased the bacterial load. Our findings indicate that IFNγ and IL-22 work in concert to accelerate production and removal of DCCs, respectively, to deny *C.r* a stable cellular substrate for colonization. Because *C.r* lacks flagella and is therefore immotile^127, 128^, its lateral spread to adjacent IECs on the colon surface from pioneering microcolonies requires host cell-to-cell migration, making accelerated clearance of infected colonocytes an effective host strategy to deny *C.r* purchase. In essence, by speeding up the “escalator” of maturing DCCs as they move up and out of the crypts and off the colonic surface, the coordinated actions of IFNγ and IL-22 restrain *C.r*’s lateral spread to prevent invasion of crypts. Notably, however, deficiency of IL-22 alone results in the invasion of crypts^4^, suggesting that despite IFNγ-driven hyperplasia of crypts, in absence of IL-22’s actions to promote accelerated shedding of *C.r*-laden DCCs—or other mechanisms driven by IL-22—crypt hyperplasia alone is ineffectual at blunting the spread of *C.r* into the crypts. Although the pathways that mediate this antagonism between pathogen-injected bacterial effectors and IL-22–induced host cell responses will require further study, the ability to isolate *C.r*-DCC conjugates for future mechanistic studies holds promise.

In a prior study, we found that IL-22–producing immune cells target different IECs within the colonic epithelium during *C.r* infection, with innate cells (predominantly ILCs) targeting surface IECs early and T cells targeting both superficial and crypt-lining IECs later to induce greater IL-22 signaling flux that correlated with protection of intestinal crypts from bacterial invasion^4^. Findings herein implicate a direct role for antigen presentation by IECs in recruiting enhanced signaling from IL-22–producing T cells as a means to specifically focus a greater cytokine payload to infected cells, whether by retaining greater numbers of IL-22-competent Th17 and Th22 cells, eliciting local secretion of IL-22 onto antigen-presenting IECs, or both. This represents another point of intersection for coordination of IFNγ and IL-22 signaling into IECs, as our results establish that IFNγ-induced up-regulation of the antigen-presenting function of IECs is required to elicit the heightened potency of T cell-derived IL-22. It also begs important new questions: in view of the restriction of *C.r* to the superficial colonocytes and near-uniform activation of crypt-lining IECs by T cell-derived IL-22, are *C.r* antigenic peptides “shared” between superficial and crypt-lining IECs?; do bacterial effector proteins injected into IECs select for TCR specificities that foster T cell-IEC interactions?; assuming that injected bacterial effectors are a major source of antigens processed and presented by IFNγ-activated IECs, is cross-presentation a major pathway by which IECs recruit T-cell help?; and, finally, is the requirement for coordination of IFNγ and IL-22 signaling a basis for the proclivity of Th17 cells to transdifferentiate into Th1-like cells in the inflamed intestines^129, 130^, i.e., does the original speculation of a “two-for-one” effector T cell lineage ascribed to Th17 reflect an evolutionary advantage in delivering for both IL-22 or IL-17 and IFNγ to most efficiently deliver Th17 help to barrier epithelia? Findings in this study provide new insights into the importance of epithelial expression of MHCII antigen presentation machinery as a mechanism by which T cells, but not ILCs, provide indispensable protection of the intestinal barrier. In addition to elucidating the basis for geographical restriction of colonization to the distal colon, these findings open a new window into our understanding of the host dynamics by which A/E enteropathogens are resisted and ultimately cleared, and will enable new studies to elucidate the competing strategies of the pathogen and host to control the survival of infected colonocytes.

## Methods

### Mice

*Il22*^hCD4.fl^ reporter/floxed and *Il22*^ΔTcell^ cKO mice were previously generated within our laboratory^4^. C57BL/6 (WT), *H2-Ab1* floxed, *mCd4-*cre and *Villin*-cre mice were purchased from Jackson Laboratory. *H2-Ab1*^Villin^ (*H2-Ab1* floxed x *Villin*-cre) cKO mice were screened by PCR and flow cytometry to exclude mice with germline deletion of MHCII^131^. All mouse strains were bred and maintained at UAB in accordance with IACUC guidelines.

### Citrobacter rodentium Infections

*Citrobacter rodentium* (*C.r*) strain, DBS100 (ATCC) was used for scRNA-seq experiments. For flow cytometry analysis and to track *C.r in situ*, we used a strain of *C.r* expressing GFP^91^ (derived from DBS100; kindly provided by Dr. Bruce A. Vallance). Mice were inoculated with 2×10^9^ cfu in a total volume of 100 µl of PBS via gastric gavage.

### Isolation of intestinal epithelial cells (IECs)

Intestinal tissue was flushed, cut into regions (5 cm ileum, 4 cm mid-distal colon or 2 cm each for proximal, middle, or distal colon), opened longitudinally and then cut into strips of 1 cm length. Tissue pieces were incubated for 20 min at 37**°**C with 1 mM DTT (Sigma), followed by 2 mM EDTA (Invitrogen) in H5H media (1x HBSS, 5% FBS, 20 mM Hepes, and 2.5 mM 2-β-ME). Tissue pieces were vortexed briefly after each 20 min incubation, followed by washing with H5H prior to centrifugation at 1800 rpm for 10 min at 4**°**C. IECs were then purified on a 40%/75% Percoll gradient by centrifugation for 20 min at 25**°**C and 2000 rpm with no brake. For analysis of *C.r*-GFP attached to IECs, tissue pieces from 4 cm of mid-distal colon were incubated for 20 min at 37**°**C with 1 mM EDTA in H5H media, followed by gentle mixing and washing with H5H.

### Single-cell (sc) RNA-sequencing and data analysis

Cells were isolated with DTT/EDTA (as described above) from mid-distal colons of naïve (BL/6), and D9 *C.r*-infected control (*Il22*^hCD4^) and T cell-specific IL-22 cKO (*Il22*^ΔTcell^) mice. Cells were pooled from 2-3 mice per condition. Live cells were sorted into 5 μl PBS with 5% FBS and 0.1 mM EDTA in an Eppendorf tube using BD Aria II. Prior to loading on the 10x Genomics Chromium instrument, cells were counted using a hemocytometer and viability of at least 90% for all samples were confirmed by trypan blue staining for a targeted number of 10,000 live cells. scRNA-Seq libraries were prepared using the Chromium Single Cell 3’ Library Kit (10x Genomics) at the FCSC Core Facility at UAB. Libraries were sequenced on an Illumina NovaSeq 6000 at the Sequencing Core Facility at La Jolla Institute, California. Cell counts were produced by Cell Ranger pipelines and transformed into Seurat objects (see Extended Data Methods Section). Data is publicly available in GEO^114^ under GSE227331.

## Acknowledgments

We thank Wanyin Deng (University of British Columbia) for helpful discussions. We gratefully acknowledge Vidya Sagar and Shanrun Liu at the UAB Flow Cytometry and Single Cell (FCSC) Core Facility, Shawn Williams at the UAB High Resolution Imaging Facility (HRIF) and J. Day at the Sequencing Core Facility at the La Jolla Institute. This work was supported by R01 grant funds from NIH/NIAID (C.T.W.) and NIH T32 support (C.G.W).

## Author Contributions

C.L.Z. and C.T.W. conceptualized the project, designed the experiments, interpreted the results, and wrote the manuscript. C.L.Z. and C.G.W. performed most of the experiments, and A.S.C. assisting with RNA work. C.G.W. and A.S.C. provided critical discussion. S.H. and Y.N. conducted *in vitro* fertilization procedures to maintain all mouse lines with the same C57BL/6J flora. R.D.H. assisted with scRNA-seq experiments. D.A.F. and M.G. carried out bioinformatics analyses of scRNA-seq data.

## Extended Data Materials and Methods

### Extended Single-cell (sc) RNA-sequencing and data analysis

#### Processing of single-cell sequencing raw data

Raw sequencing data were processed with the Cell Ranger pipeline software (v.3.0.2; 10x Genomics). Raw base call files generated by Novaseq 6000 (Illumina) were converted to FASTQ files using *cellranger mkfastq* with default parameters. The Cell Ranger count pipeline was used to perform quality control, sample demultiplexing, barcode processing, alignment, and single-cell 5’ gene counting. Cell ranger “count” was used to align raw reads against the GRCm38 mouse reference transcriptome. Subsequently, cell barcodes and unique molecular identifiers (UMI) underwent filtering and correction using default parameters in Cell Ranger. Reads with the retained barcodes were quantified and used to build the gene expression matrix.

#### Classification of IECs using single-cell transcriptome data

Seurat^132^ (v.3.0.0), implemented using the R package, was applied to exclude low-quality cells. Cells that expressed fewer than 200 genes were filtered out. Genes that were not detected in at least three single cells were excluded. EpCAM^+^ cells were selected and based on these criteria, we retained the total numbers of IECs per genotype: 5064 cells from naïve BL/6, 6081 cells from d9 *C.r*-infected *Il22*^hCD4^ (control), and 4416 cells from d9 *C.r*-infected *Il22*^ΔTcell^ pooled samples. The processed data was normalized using Seurat’s ‘NormalizeData’ function, which used a global scaling normalization method, LogNormalize, to normalize the gene expression measurements for each cell to the total gene expression. Highly variable genes were then identified using the function ‘FindVariableGenes’ in Seurat. The anchors were identified using the ‘FindIntegrationAnchors’ function, and thus the matrices from different samples were integrated with the ‘IntegrateData’ function. The variation arising from library size and percentage of mitochondrial genes was regressed out using the function ‘ScaleData’ in Seurat. Principal component (PC) analysis was performed using the Seurat function ‘RunPCA’, and K-nearest neighbor graph was constructed using ‘FindNeighbors’ function in Seurat with the number of significant PCs identified from PCA analysis. Clusters were identified using ‘FindClusters’ function with resolution of 0.8. The clusters were visualized in two dimensions with UMAP. The normalization, integration, and clustering were performed under standard Seurat workflow.

#### Identification of differentially expressed genes

Differential gene expression analyses were carried out using the Seurat function ‘FindMarkers’. Briefly, we performed the Wilcoxon rank-sum test with the default threshold of 0.25 for log2 fold change and a filter for the minimum percent of cells in a cluster greater than 25%. The differentially expressed genes were isolated by comparing significantly upregulated genes and downregulated genes defined as adjusted P-value, Padj < 0.05. Top differentially expressed genes were visualized by heatmap via ComplexHeatmap (v2.11.1) package in R. Expression of individual differentially expressed genes were represented as violin plots. Violin plots were rendered using the function ‘VlnPlot’ in Seurat.

#### Gene Set Enrichment Analysis (GSEA)

The fgsea R package^133^ (v1.4.0) was used for gene set enrichment. Gene sets used were Gene Ontology Biological Process (GO_BP) gene sets. The input for GSEA was a set of gene signatures of a cluster or cell type obtained from Seurat. A p-value quantifying the likelihood that a given gene set displays the observed level of enrichment for genes was calculated using Fast Gene Set Enrichment Analysis (fgsea, v1.4.0) with 1 million permutations. Gene set enrichment p-values of Normalized Enrichment Scores (NES) were corrected with the Benjamini-Hochberg procedure^134^. The top enriched terms were visualized with dot plots using R package ggplot2 (v3.3.5).

#### Trajectory analysis

RNA velocity analysis was conducted using the scVelo package (v0.2.2) with Scanpy (v1.6.1) on Python (v3.8.5). To perform trajectory analysis, the un-spliced and spliced variant count matrix that was calculated using 10× pipeline in velocyto (v0.17.16) was fused with an anndata object containing the UMAP information and cluster identities defined in Seurat analysis. The combined dataset was then processed using the scVelo pipeline: The ratio of un-spliced:spliced RNA for each gene was filtered and normalized using the default settings. Afterward, the first and second moments were calculated for velocity estimation. Following moment calculation, the dynamic model was used to calculate the RNA velocities. The dynamic model iteratively estimates the parameters that best model the phase trajectory of each gene, therefore capturing the most accurate, albeit more computationally intensive, estimate of the dynamics for each gene. These approaches were used to graphically model the RNA velocity for each condition.

#### Data availability

The scRNA-seq data used in this study have been deposited in the Gene Expression Omnibus database under the accession number GSE227331. All other data generated in this study are provided within the article and its Supplementary Information/Source Data file. The shell, R and Python scripts that enabled the main steps of the analyses performed in this project are available on request.

### Tissue Preparation

Tissues were fixed in cold 4% PFA overnight at 4**°**C. Next day, tissue was rinsed in cold 1x PBS for several washes including an overnight incubation at 4**°**C. Next day, tissue was placed in cold 30% sucrose in 1x PBS overnight at 4**°**C. Tissue was embedded in O.C.T. (Tissue-Tek) and frozen with 2-methyl butane chilled with liquid nitrogen. For pSTAT3 staining, tissue sections were permeabilized in cold methanol (Fisher) for 10-15 min at −20**°**C. Tissue sections were blocked at RT for 30 minutes with 10% mouse serum in 1x PBS and 0.05% Tween-20. Antibodies were diluted in 2% BSA/PBS/Tween-20 and incubated for 20-30 min at RT or ON’ at 4**°**C (pSTAT3 stain). The following antibodies/reagents were used: anti-BrdU (BU1/75; Abcam), anti-EpCAM1 (G8.8; ThermoFisher), anti-Fabp2 (polyclonal; R&D/Fisher), anti-GFP (A-11122; ThermoFisher), anti-goat IgG (ThermoFisher), anti-Ly6g (1A8; Biolegend or ThermoFisher), anti-rabbit IgG (ThermoFisher), Streptavidin-Alexa Fluor 594, UEA-1 (Vector Labs) and Prolong Diamond antifade mountant with DAPI (ThermoFisher).

### Flow Cytometry

Colonic IECs were stained with Fc Block (Clone 2.4G2) followed by staining with fluorescent-labeled antibodies in IEC buffer (1x PBS with 5% FBS and 2mM EDTA to reduce cell clumping) on ice in 1.5 ml microcentrifuge tubes. For intracellular staining, cells were fixed and permeabilized using BD Cytofix/Cytoperm kit (BD Bioscience). Samples were acquired on an Attune NxT flow cytometer (Life Technologies) and analyzed with FlowJo software. Cells were sorted on either a BD FACS Aria or Aria II (BD Biosciences). The following antibodies/reagents were used: anti-CD45 (30-F11; Biolegend), anti-EpCAM1 (G8.8; ThermoFisher), anti-Fabp2 (polyclonal; R&D/Fisher), anti-goat IgG (ThermoFisher), anti-Ly6G (1A8; Biolegend), anti-MHCII (M5/114.15.2; Fisher) and Live/Dead Fixable Near-IR dead cell dye (ThermoFisher).

### Real Time PCR

cDNA synthesis was performed with iScript reverse transcription (RT) Supermix (Bio-Rad) according to manufacturer’s instructions. cDNA amplification was analyzed with SsoAdvanced Universal SYBR Green Supermix (Bio-Rad) in a Biorad CFX qPCR instrument. See primer list in Extended Data Table 2.

### Colony counts of *C.r*-GFP

Feces were collected, weighed, and dispersed for 30 seconds in 1x PBS using a PowerGen500 homogenizer (Fisher). Liver was removed under sterile conditions, placed in 2-3 ml H5H in Miltenyi M tubes, weighed, and homogenized using Miltenyi GentleMACS Dissociator using Program RNA_01. Homogenate was filtered through a 70 μm cell strainer and then spun at 8000 rpm for 15 minutes to pellet cells. Cell pellet was resuspended in 1x PBS and serially diluted and plated in duplicate on LB plates containing 30 μg/ml Chloramphenicol. Colonies were counted after 24 hr incubation at 37°C to determine the log10 CFU per gram of tissue.

