## Supplemental Data for "A novel subset of colonocytes targeted by *Citrobacter rodentium* elicits epithelial MHCII-restricted help from CD4 T cells"

### Supplemental Figure Legends

**Supplemental Data Fig. 1 Characterization of colonocytes from naïve BL/6 mice.** scRNA-seq was performed on colonic epithelial cells isolated from mid-distal colon of naïve BL/6 mice. **a**, UMAPs and **b**, Violin plots of common enterocyte markers. **c**, Dot plot of glycoproteins expressed by IECs from naïve BL/6 mice. **d**, UMAPs and **e**, Violin plots of DCC lineage markers. **f**, UMAPs and **g**, Violin plots of PCC lineage markers. 2 mice pooled per sample, *n*=2 biological replicates per group. IEC=intestinal epithelial cell; CC=colonocyte; PCC=middle colonocyte; DCC=distal colonocyte; DCS=deep crypt secretory cell; TA=transit-amplifying cell; CBC=crypt base columnar cell; Undiff=undifferentiated cell. *Car4* (carbonic anhydrase 4) and *Slc26a3* (chloride ion transporter) are strongly expressed by both PCC and DCC lineage colonocytes but not CBC cells, progenitor cells, or secretory IECs. Top genes expressed by naïve mature DCCs include *Ly6g*, *Slc20a1*, and *Dmbt1*. The top genes expressed by naïve pre- and mature PCCs include *Fabp2*, *Dpep1*, and *Emp1*. Interestingly, PCC and DCC lineage colonocytes express different glycoproteins which may dictate differences in cellular communication and interaction with specific molecules or neighboring cells.

**Supplemental Data Fig. 2 *C.r* infection causes enhanced IEC proliferation in the crypts. a**, Colon tissue was collected at various times after d9 *C.r*-infected BL/6 mice were treated with BrdU and stained for BrdU (red), EpCAM1 (green) and DAPI (blue). Scale bar, 50 μm. **b**, Percent migration of BrdU$^{+}$ IECs from the crypt base to the luminal surface per total length of the crypt. 4 mice per time point per group, 20-30 crypts per group, *n*=1. One-way ANOVA; ***p≤0.001 comparing naïve and d9 *C.r*-infected mice. Data are represented as mean ± SEM. **c**, Dot plots of top markers for all populations from scRNA-seq analysis of IECs from naïve and d9 *C.r*-infected mice. 2 mice pooled per sample, *n*=2 biological replicates per group. IEC=intestinal epithelial cell; CC=colonocyte; PCC=middle colonocyte; DCC=distal colonocyte; DCS=deep crypt secretory cell; TA=transit-amplifying cell; CBC=crypt base columnar cell.

**Supplemental Data Fig. 3 CD4 T cells upregulate IL-22–inducible genes on colonocytes and goblet cells for host defense. a-b**, Mid-distal colon IECs from d8 *C.r*-GFP-infected mice were isolated and stained for EpCAM1, Ly6G, and CD45 and either **a**) analyzed by flow cytometry or **b**) sorted on EpCAM1$^{+}$ (red) *C.r*-GFP$^{+}$ (green) cells, cytospun, and stained with DAPI (blue). Scale bar = 10 μm. 3 mice per group and *n*=2 independent experiments. **c**, IECs from naïve (open symbol) and d8 *C.r*-infected mice (closed symbol) were sorted from distal ileum (green), proximal colon (blue) and distal colon (red) and *Lcn2*, *Lbp*, *Cxcl2* and *Muc1* mRNA expression was analyzed by RT-PCR. 2-3 mice pooled per sample seq and *n*=2 independent experiments. One-way ANOVA with Tukey's multiple comparison test; ***p≤0.001 comparing naïve and infected mice; and $^{øøø}$p≤0.001 comparing infected samples from different tissue regions. Data are represented as mean ± SEM. **d-e**, scRNA-seq was performed on colonic epithelial cells isolated from mid-distal colon of naïve BL/6, and d9 *C.r*-infected *Il22*$^{hCD4}$ (Control) and *Il22*$^{\Delta Tcell}$ cKO mice. Dot plots of fucosyl transferases and mucins (**d**) and IL-22-inducible genes (**e**) from naïve, and d9 *C.r*-infected *Il22*$^{hCD4}$ and *Il22*$^{\Delta Tcell}$ mice. 2 mice pooled per sample, *n*=2 biological replicates per group. IEC=intestinal epithelial cell; CC=colonocyte; PCC=middle colonocyte; DCC=distal colonocyte; DCS=deep crypt secretory cell; TA=transit-amplifying cell; CBC=crypt base columnar cell. As expected in naïve mice, goblet cells predominantly express *Muc2* and *Muc4*; whereas colonocytes express *Muc3* and *Muc13*. *Fut2* gene which encodes for the fucosyl transferase 2 enzyme is upregulated on both goblet cells and colonocytes; whereas *Muc1*, a membrane-bound mucin is predominantly upregulated on stem cells and colonocytes of the DCC lineage during *C.r* infection. *S100a8/9* antimicrobial peptide, and *Cxcl2* and *Cxcl5 chemokines* are predominantly expressed by mature colonocytes of the DCC lineage and are heightened in the presence of IL-22-producing CD4 T cells suggesting they may play important roles in bacterial clearance during the late phase of *C.r* infection.

**Supplemental Data Fig. 4 CD4 T cells upregulate IL-22–inducible genes on DCC lineage cells.** scRNA-seq was performed on colonic epithelial cells isolated from mid-distal colon of naïve

BL/6, and d9 *C.r*-infected *Il22*$^{hCD4}$ (Control) and *Il22*$^{\Delta Tcell}$ cKO mice. Violin plots of *S100a8*, *Cxcl5* and *Lrg1* expression in pre-DCC (beige), pro-DCC (orange), mature DCC (royal blue) and pathogen-induced (P-I) CC from naïve, and *C.r* d9 *Il22*$^{fl/fl}$ and *Il22*$^{\Delta Tcell}$ mice. 2 mice pooled per sample, *n*=2 biological replicates per group. Bars represent mean gene expression and dots denote gene expression per individual cell.

**Supplemental Data Fig. 5 CD4 T cells upregulate IL-22–inducible genes on multiple colonic IEC subsets.** scRNA-seq was performed on colonic epithelial cells isolated from mid-distal colon of naïve BL/6, and d9 *C.r*-infected *Il22*$^{hCD4}$ (Control) and *Il22*$^{\Delta Tcell}$ cKO mice. **a**, UMAPs and **b**, Violin plots from naïve BL/6 mice. **c**, UMAPs and **d**, Violin plots from *C.r* d9 Control mice. **e**, UMAPs and **f**, Violin plots from *C.r* d9 *Il22*$^{\Delta Tcell}$ mice. 2 mice pooled per sample, *n*=2 biological replicates per group. *S100a* family of AMPs and neutrophil-recruiting chemokines (e.g., *Cxcl2* and *Cxcl5*) are predominantly upregulated on IECs of the mature DCC lineage in infected control mice and expression of these genes is severely blunted in the absence of IL-22–producing T cells. *Lrg1*, a leucine-rich α-2-glycoprotein thought to play a role in cell migration and wound healing[105,106] is upregulated on all IECs during *C.r* infection. IL-22–producing T cells augment Lrg1 expression on most IECs with the greatest expression observed on pre-DCCs and pro-DCCs suggesting that T cell-driven upregulation of *Lrg1* may contribute to the expansion of distal colonocytes and ultimate restitution of the damaged epithelium during *C.r* infection. *Fut2*, another known IL-22–regulated gene[22] is upregulated on goblet cells and colonocytes of the DCC lineage in response to IL-22$^{+}$ T cells on d9 of *C.r* infection.

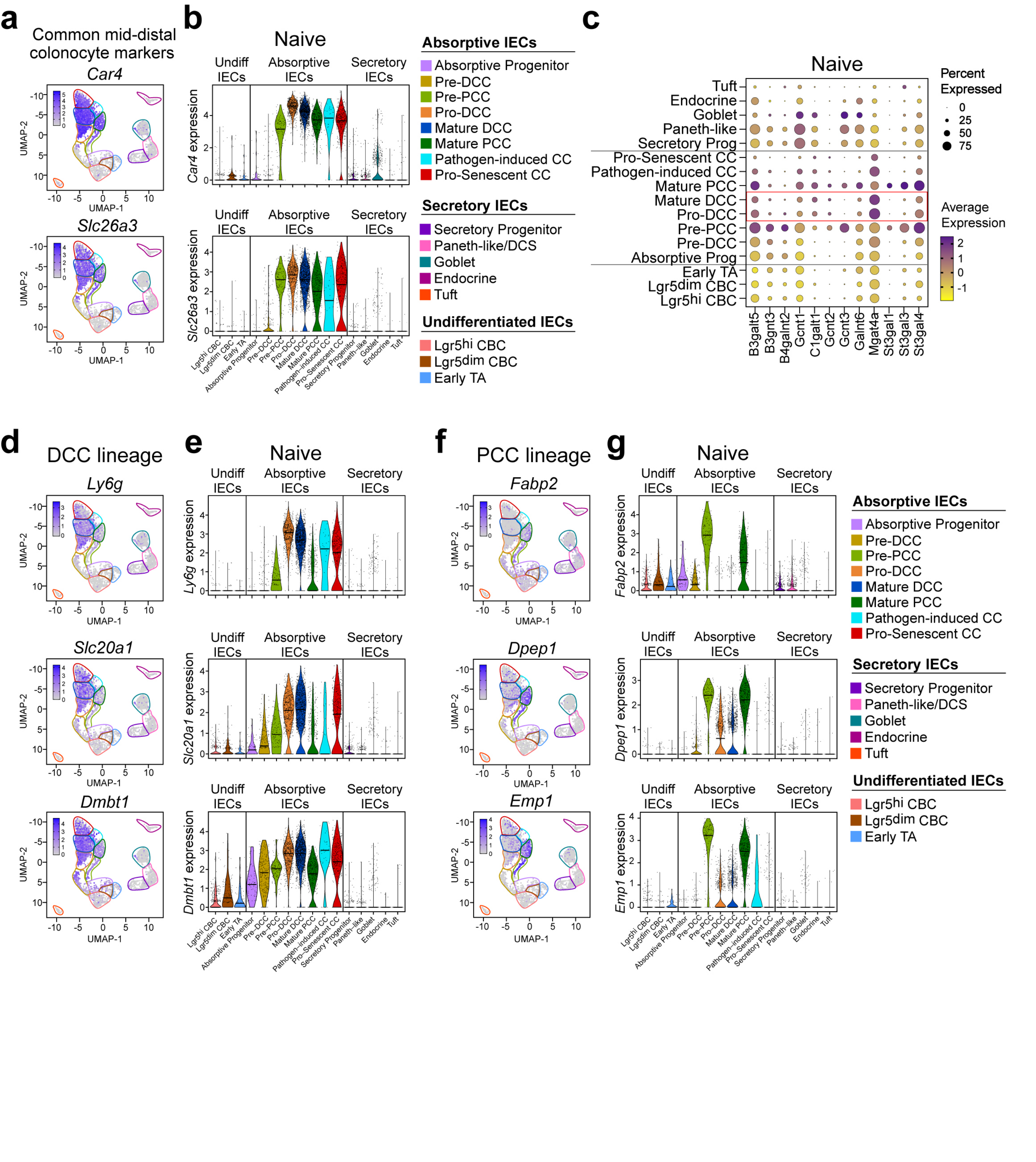
a
Common mid-distal colonocyte markers
Car4
Slc26a3
UMAP-1
UMAP-2
b
Naive
Undiff IECs
Absorptive IECs
Secretory IECs
Car4 expression
Slc26a3 expression
Absorptive IECs
Absorptive Progenitor
Pre-DCC
Pre-PCC
Pro-DCC
Mature DCC
Mature PCC
Pathogen-induced CC
Pro-Senescent CC
Secretory IECs
Secretory Progenitor
Paneth-like/DCS
Goblet
Endocrine
Tuft
Undifferentiated IECs
Lgr5hi CBC
Lgr5dim CBC
Early TA
c
Naive
Tuft
Endocrine
Goblet
Paneth-like
Secretory Prog
Pro-Senescent CC
Pathogen-induced CC
Mature PCC
Mature DCC
Pro-DCC
Pre-PCC
Pre-DCC
Absorptive Prog
Early TA
Lgr5dim CBC
Lgr5hi CBC
B3galt5
B3gnt3
B4galnt2
Gcnt1
C1galt1
Gcnt2
Gcnt3
Galnt6
Mgat4a
St3gal1
St3gal3
St3gal4
Percent Expressed
0
25
50
75
Average Expression
2
1
0
-1
d
DCC lineage
Ly6g
Slc20a1
Dmbt1
e
Naive
Ly6g expression
Slc20a1 expression
Dmbt1 expression
f
PCC lineage
Fabp2
Dpep1
Emp1
g
Naive
Fabp2 expression
Dpep1 expression
Emp1 expression

ED Figure 1

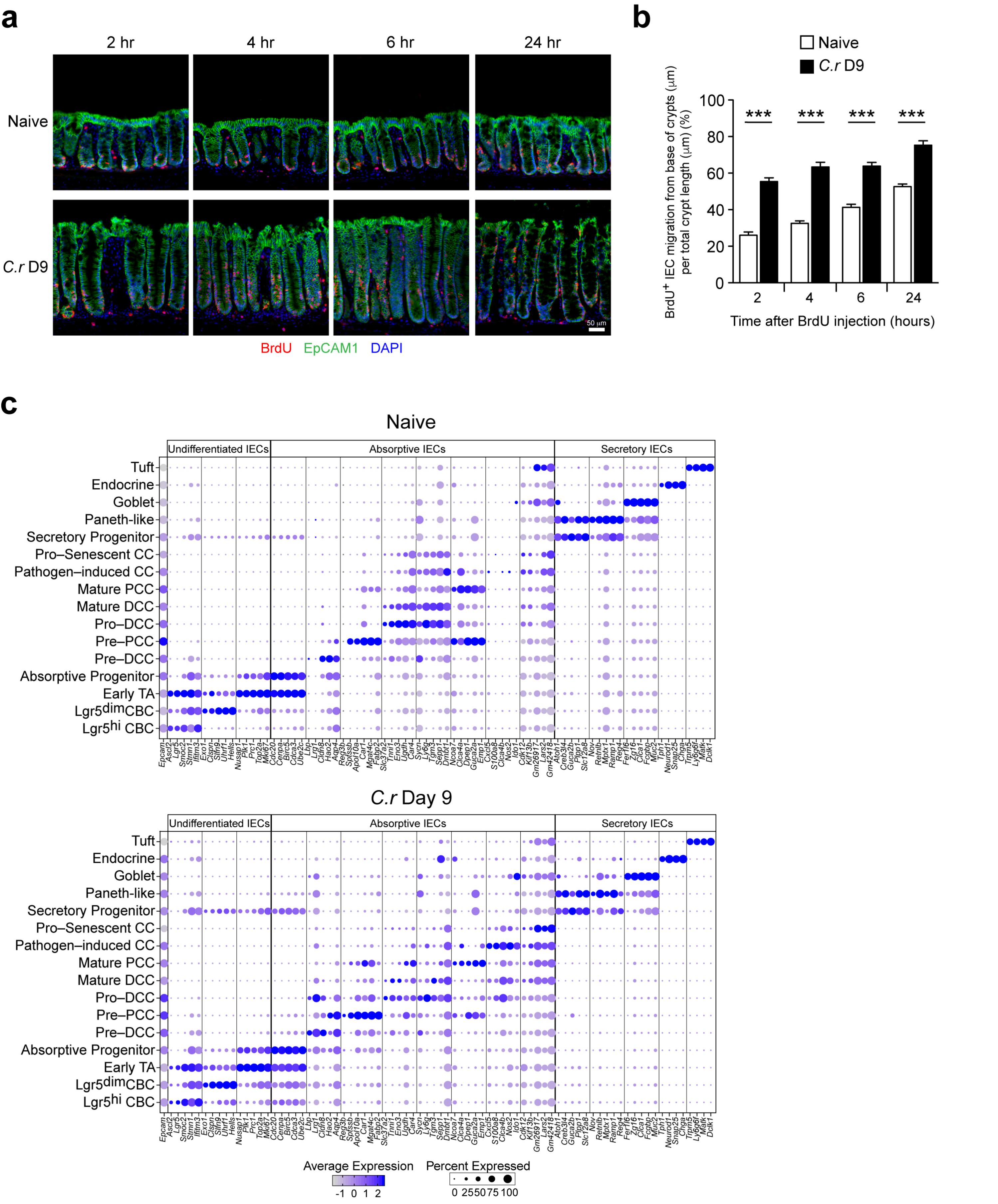
a
2 hr
4 hr
6 hr
24 hr
Naive
C.r D9
50 µm
BrdU EpCAM1 DAPI
b
Naive
C.r D9
BrdU+ IEC migration from base of crypts (µm) per total crypt length (µm) (%)
***
Time after BrdU injection (hours)
c
Naive
Undifferentiated IECs
Absorptive IECs
Secretory IECs
Tuft
Endocrine
Goblet
Paneth–like
Secretory Progenitor
Pro–Senescent CC
Pathogen–induced CC
Mature PCC
Mature DCC
Pro–DCC
Pre–PCC
Pre–DCC
Absorptive Progenitor
Early TA
Lgr5dimCBC
Lgr5hi CBC
C.r Day 9
Average Expression
Percent Expressed
0 25 50 75 100

ED Figure 2

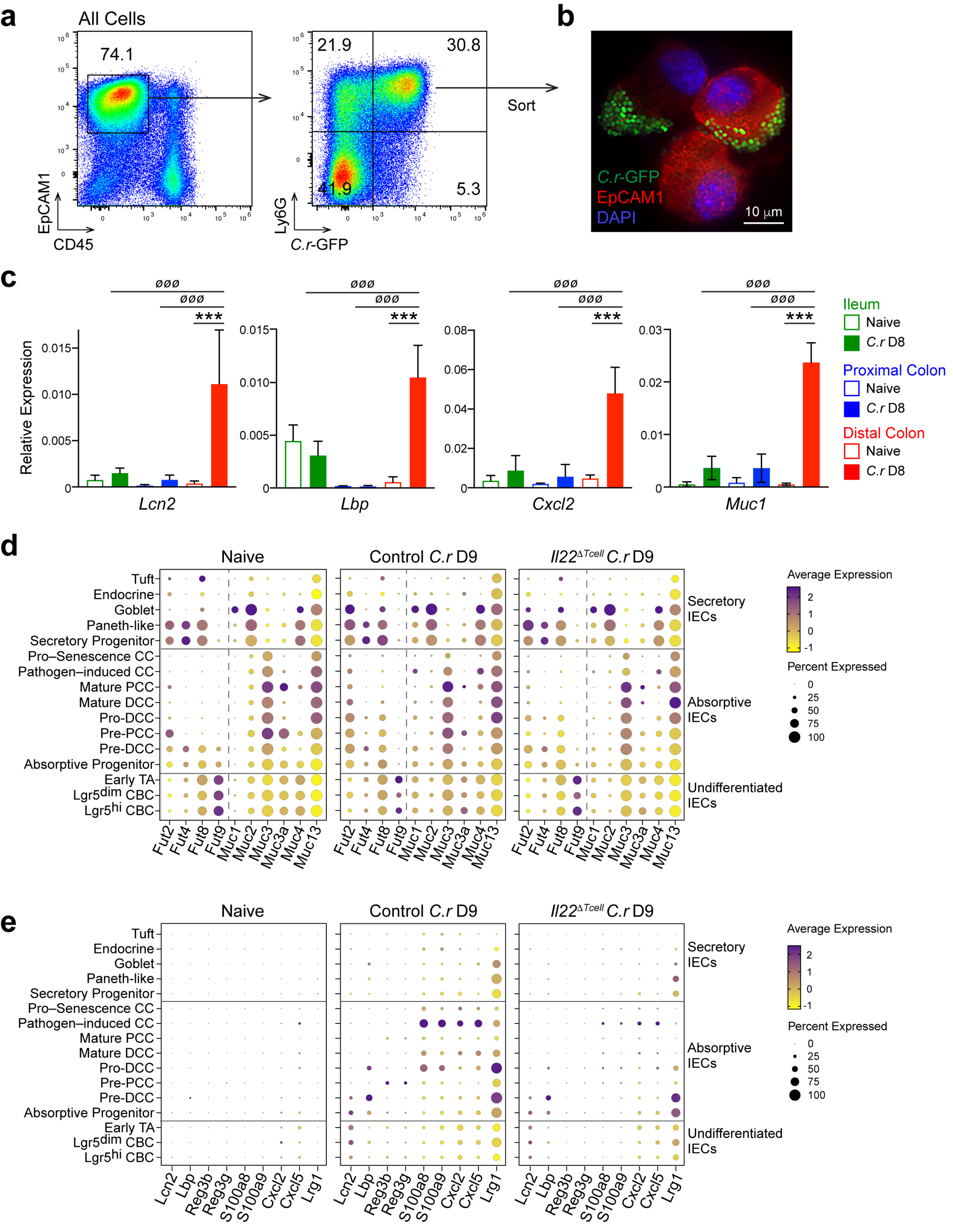
a
All Cells
74.1
EpCAM1
CD45
21.9
30.8
41.9
5.3
Ly6G
C.r-GFP
Sort
b
C.r-GFP
EpCAM1
DAPI
10 μm
c
Relative Expression
Lcn2
Lbp
Cxcl2
Muc1
Ileum
Naive
C.r D8
Proximal Colon
Naive
C.r D8
Distal Colon
Naive
C.r D8
d
Naive
Control C.r D9
Il22ΔTcell C.r D9
Tuft
Endocrine
Goblet
Paneth-like
Secretory Progenitor
Pro–Senescence CC
Pathogen–induced CC
Mature PCC
Mature DCC
Pro-DCC
Pre-PCC
Pre-DCC
Absorptive Progenitor
Early TA
Lgr5dim CBC
Lgr5hi CBC
Fut2 Fut4 Fut8 Fut9 Muc1 Muc2 Muc3 Muc3a Muc4 Muc13
Secretory IECs
Absorptive IECs
Undifferentiated IECs
Average Expression
Percent Expressed
e
Naive
Control C.r D9
Il22ΔTcell C.r D9
Lcn2 Lbp Reg3b Reg3g S100a8 S100a9 Cxcl2 Cxcl5 Lrg1
Secretory IECs
Absorptive IECs
Undifferentiated IECs
Average Expression
Percent Expressed

**ED Figure 3**

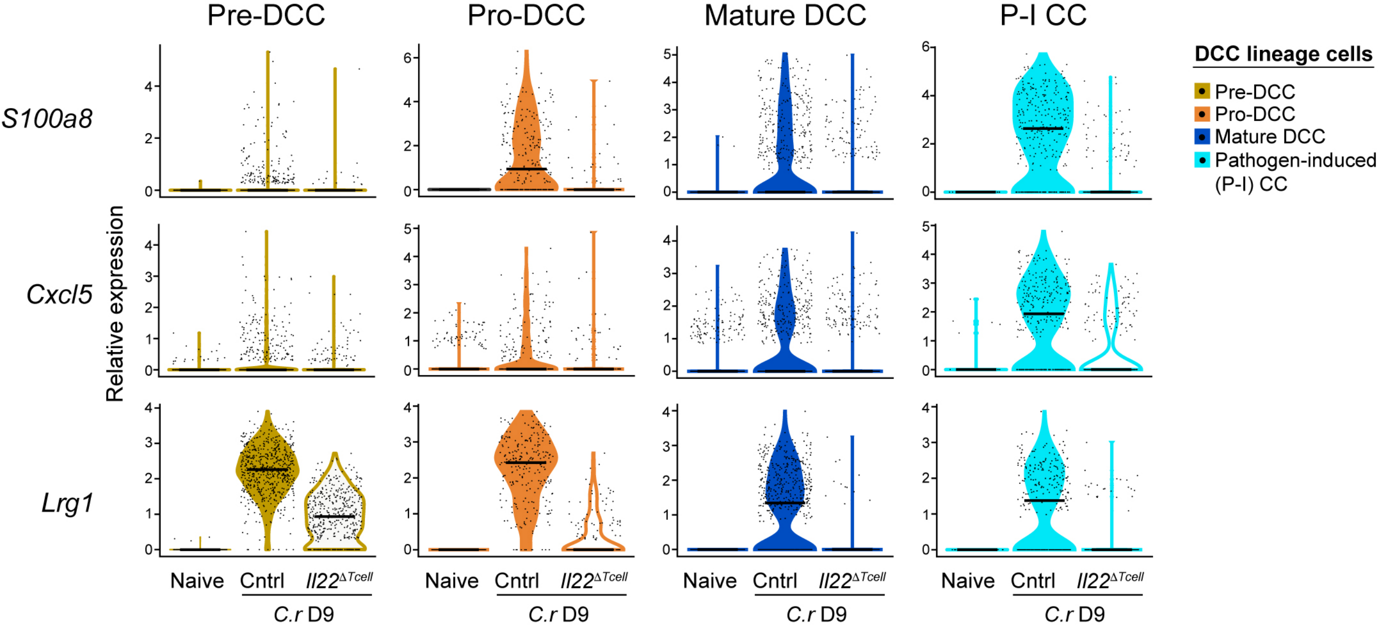

Pre-DCC
Pro-DCC
Mature DCC
P-I CC
DCC lineage cells
Pre-DCC
Pro-DCC
Mature DCC
Pathogen-induced (P-I) CC
S100a8
Cxcl5
Lrg1
Relative expression
Naive
Cntrl
Il22ΔTcell
C.r D9

**ED Figure 4**

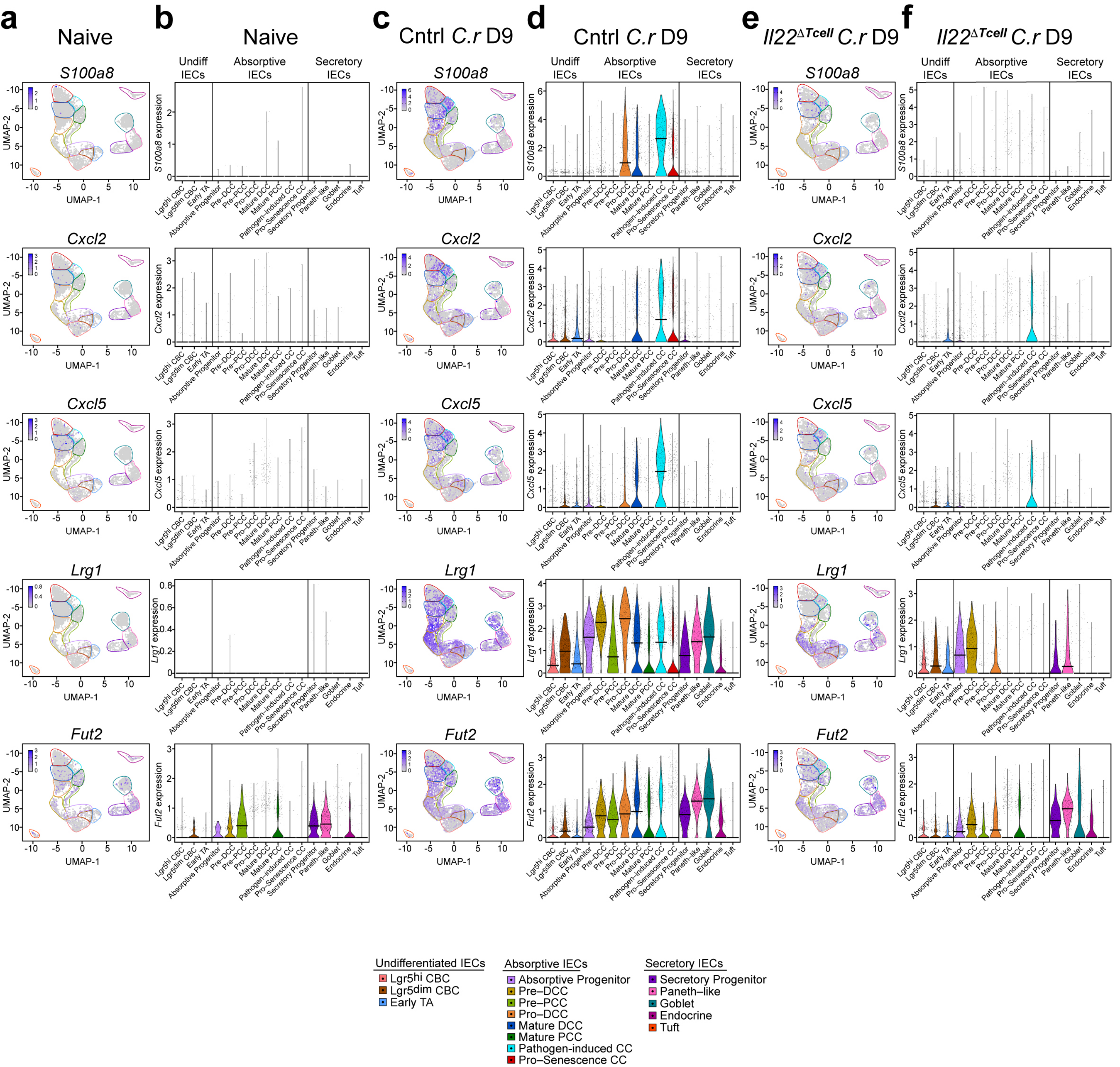
a Naive
b Naive
c Cntrl C.r D9
d Cntrl C.r D9
e Il22ΔTcell C.r D9
f Il22ΔTcell C.r D9
Undiff IECs
Absorptive IECs
Secretory IECs
S100a8
Cxcl2
Cxcl5
Lrg1
Fut2
UMAP-1
UMAP-2
S100a8 expression
Cxcl2 expression
Cxcl5 expression
Lrg1 expression
Fut2 expression
Lgr5hi CBC
Lgr5dim CBC
Early TA
Absorptive Progenitor
Pre–DCC
Pre–PCC
Pro–DCC
Mature DCC
Mature PCC
Pathogen–induced CC
Pro–Senescence CC
Secretory Progenitor
Paneth–like
Goblet
Endocrine
Tuft
Undifferentiated IECs
Absorptive IECs
Secretory IECs

ED Figure 5

| Lgr5-hi | Lgr5-dim | Early TA cells | Absorptive Progenitor | Pre-DCC | Pre-MCC | Pro-DCC | Mature DCC | Mature MCC | Pathogen-induced CC | Pro-Senescent CC | Secretory Progenitor | Paneth-like cells | Goblet cells | EEC | Tuft |
|---|---|---|---|---|---|---|---|---|---|---|---|---|---|---|---|
| Lgr5 | Hells | Nusap1 | Ube2c | Ly6a | Aqp8 | Tnni1 | Car4 | Emp1 | S100a8 | Lars2 | Atoh1 | Agr2 | Zg16 | Chga | Matk |
| Smoc2 | Slfn9 | Kif11 | Cdc20 | Lrg1 | Fabp2 | Slc37a2 | Dmbt1 | Dpep1 | Ido1 | Atp12a | Txndc5 | Spink4 | Clca1 | Tph1 | Dclk1 |
| Rgcc | Uhrf1 | Kif20b | Birc5 | Ly6c1 | Sptssb | Fxyd4 | Tgm3 | Guca2a | Ubd | Tnfaip3 | Guca2b | Reg4 | Fcgbp | Neurod1 | Trpm5 |
| Trf | Clspn | Prc1 | Tpx2 | Lbp | Reg3b | Sycn | Ly6g | Ndrg1 | Cxcl2 | Irf1 | Dll1 | Mptx1 | Tff3 | Snap25 | Ly6g6f |
| Acot1 | Exo1 | Cdk1 | Cenpa | Cldn8 | Reg3g | Saa1 | Slc20a1 | Ncoa7 | Cxcl5 | Kif13b | Cbfa2t3 | Retnlb | Muc2 | Pyy | Dnah5 |
| Ascl2 | Tcf19 | Plk1 | Cdkn3 | Hao2 | Apol10a | Ugdh | Slc26a3 | Cyp2c55 | Nos2 | Tle4 | Plpp1 | Ramp1 | Spink1 | Sct | Alox5 |
| Cdca7 | Top2a | Mki67 | Cdca3 | Gsdmc4 | Higd1a | Eno3 | Cyp2d9 | Clca4a | Xdh | Mprip | Slc12a8 | Nov | Fer1l6 | Pcsk1n | |
| Stmn1 | | | | Aqp4 | Mgat4c | | Sepp1 | Muc3 | Gbp2 | Arap2 | Creb3l4 | | | | |
| Ifitm3 | | | | | Car1 | | | | Tgm2 | Cdk12 | | | | | |
| | | | | | | | | | Tnf | Myo15b | | | | | |
| | | | | | | | | | Cd274 | Gm26917 | | | | | |
| | | | | | | | | | Adrb2 | Gm42418 | | | | | |
| | | | | | | | | | Clca4b | | | | | | |
| | | | | | | | | | Upp1 | | | | | | |

| Gene name | Forward Primer | Reverse Primer |
|---|---|---|
| *Ly6g* | GAC TTC CTG CAA CAC AAC TA | TCA CGT TGA CAG CAT TAC C |
| *Slc20a1* | TTC CCA TCA GCA CAA CAC | CCA GTC AAC AGC CTT CTT T |
| *Slc26a3* | GCG TGT ACT CCC TCA AAT AC | GCT CCC TGC AAA TCC TTT |
| *Fabp2* | CGG TGT AAA CTT TCC CTA CAG | TGG CCT CAA CTC CTT CAT A |
| *Dpep1* | ACA AAG ATG CCG TGA AGA G | CCA AGG CTG CTG TCA ATT A |
| *Guca2a* | GCA CCA CAG CTA TGT AGT AG | GAG AAA GGC AAG CGA TGT |
| *Reg3b* | ATG GCT CCT ACT GCT CC | GTG TCC TCC AGG CCT CTT T |
| *S100a8* | TGA GTG TCC TCA GTT TGT GCA G | TGT GAG ATG CCA CAC CCA CTT T |
| *Cxcl2* | GCT GTC AAT GCC TGA AGA | TTC AGG GTC AAG GCA AAC |
| *Cxcl5* | TCT TGT CCA CAA TGA GCC TCC A | AAC AGC AAC AGA AAT GCC AGC G |
| *Muc1* | GCA TAA GAA GGA GGC AGA TG | GGG CAA GGA AAT AGA CGA TAG |
| *Fut2* | CCC ACT TCC TCA TCT TTG TC | CGC CTG TAA TTC CTT CTC TG |
| *Il22ra1* | CAC CGT CTA CAG TGT GGA ATA TAA | CGT GAC CTT GGC GTA GTA AA |
| *Lbp* | CAG CCG CAT TTG TGA TTT G | TGG CAG AGT CTG GAG ATA AG |
| *Lcn2* | TTT CAC CCG CTT TGC CAA GTC T | CAC ACT CAC CAC CCA TTC AGT TGT |
| *Lrg1* | CCT CAA GGA ATG CCT GAT AC | GAG AAT TCC ACC GAC AGA TG |
| *Eae* | CCA AAG GAA TCG GAG TGT AGT T | TAG GTG GCA AGC TGA TGT ATG |
| *Saa1* | ATT GCT GAC CAG GAA GCC AAC A | AGG ACG CTC AGT ATT TGT CAG GCA |
| *Gapdh* | TCC ATG ACA ACT TTG GCA TTG | CAG TCT GGG TGG CAG TGA |
| *Slc13a2* | GTG CTA TGT CTG CCC ATT T | CTT GGA GGC TTC CAT GAT AC |
| *Lrg1 2010* | TCC AGC CTC AAG GAA TGC CTG ATA | ATT CCA CCG ACA GAT GGA CAG T |
